# Predicting protein folding pathway using a novel folding force field model derived from known protein universe

**DOI:** 10.1101/2023.11.17.567532

**Authors:** Kailong Zhao, Pengxin Zhao, Suhui Wang, Yuhao Xia, Guijun Zhang

**Affiliations:** College of Information Engineering, Zhejiang University of Technology, HangZhou 310023, China

**Keywords:** protein folding pathway, folding force field, evolutionary history

## Abstract

The protein folding problem has emerged as a new challenge with the significant advances in deep learning driven protein structure prediction methods. While the structures of almost all known proteins have been successfully predicted, the process by which they fold remains an enigma. Understanding the intricate folding mechanism is of paramount importance, as it directly impacts the stable expression and biological function of proteins. Here, we propose FoldPAthreader, a protein folding pathway prediction method that designs a novel folding force field model by exploring the intrinsic relationship between protein evolutionary history and folding mechanisms from the known protein universe. Further, the folding force field is used to guide Monte Carlo conformational sampling, driving the protein chain fold into its native state by exploring a series of transition states and potential intermediates. On the 30 targets we collected, FoldPAthreader can successfully predict 70% of the proteins whose folding pathway is consistent with wet-lab experimental data. The results show that the folding force field can capture key dynamic features of hydrogen bonding and hydrophobic interactions. Importantly, for the widely studied BPTI and TIM proteins, the folding pathway predicted by FoldPAthreader have the same microscopic dynamic properties as those simulated by molecular dynamics.

**Significance Statement:** Protein folding is the process by which a protein acquires its functional conformations by gradually transforming from random coils into a specific three-dimensional structure. In the post-Alphafold2 era, functional analysis of protein macromolecules should not only rely on the final state structure, but should pay more attention to the structural folding process, that is, the various intermediate states formed during the folding process. At present, there is no folding force field specifically used for protein folding pathway prediction in computational biology. Here we extracted folding information from 100-million-level structure database and designed a new folding force field for folding pathway prediction, proving a hypothesis that the protein evolutionary history implicitly contains folding information of individual protein. This study may provide new insights into the understanding of protein folding mechanisms, which is expected to advance drug discovery.

## Introduction

The folding mechanism of proteins reveals fundamental processes of life^1^. Proper folding typically results in proteins existing in a soluble form within cells. If the folding rate is too slow or there are errors in the folding, it may cause the protein to exist in an insoluble form, leading to loss of protein function and even cause some diseases related to abnormal protein aggregation^2^. With knowledge of protein folding, researchers can target specific steps in the folding process to design drugs that stabilize or disrupt specific conformations to achieve the desired therapeutic effect^3^. Therefore, understanding the protein folding mechanism is of great significance for unraveling disease mechanisms and personalized medicine^4, 5^.

Protein folding is an extremely intricate process that entails the spontaneous arrangement of amino acid chains into their biologically active three-dimensional structures through a series of conformational changes. Each of these changes is influenced by the surrounding solvent context and sequence mutations^6, 7^. This complexity presents significant challenges for experimental scientists when investigating the protein folding mechanism. Researchers often employ a multi-faceted approach that combines multiple experimental techniques to obtain protein folding information from different perspectives to understand its dynamic process and the formation of intermediate states^8, 9^. The complexity of experimental techniques has driven scientists to rely on computational techniques to study protein folding mechanisms^10, 11^. Molecular dynamics (MD) is one of the popular tools for studying protein folding dynamics^12^. Unfortunately, tracking the folding process at the level of thermally driven residue-level dynamics is computationally demanding and often unfeasible for long timescales^13^, and the molecular mechanics force fields used in MD simulations are not sufficiently accurate. To overcome the time scale limitations of MD simulations and effectively explore the complex energy landscape of proteins, various efficient and enhanced sampling methods such as Pathfinder^14^, MELD^15, 16^, DBFOLD^17^ and P3Fold^18^ have been developed to study folding mechanisms^19, 20^.

However, force field models in molecular dynamics simulations or Monte Carlo (MC) conformational sampling methods typically focus on capturing stable conformations and final structures of proteins. These force fields include physical potential terms such as hydrogen bonds and hydrophobic interactions, as well as statistical potential terms like Ramachandran, to guide proteins to accurately fold into a three-dimensional structure^21^, without focusing on the topological plausibility of transition states or intermediates during the folding process^22^. Therefore, designing dedicated folding force field models specifically for predicting folding pathway and intermediates is an urgent challenge in the post-AlphaFold2 era^23^.

During early evolution, there may have been many disordered polypeptides or polypeptide-like molecules^24^. These peptides may function in their disordered structure without specific folding. As biological systems become more complex, a need may arise for specific 3D structures that can more efficiently perform certain biological functions^25^. During this process, evolutionary selection on foldable sequences may have led to the development of folding ability. The appearance and evolution of foldable sequences gradually became the basis of protein folding^26^. Therefore, we can try to establish a link between the folding process of proteins and the evolution of structure. As Ernst Haeckel claimed that ontogeny recapitulates phylogeny. He argues that individuals undergo a series of morphological changes during development that reflect the stages that species has gone through in its evolutionary history^27^. Applying this point to the study of protein folding, we can get an interesting hypothesis that the conformational changes that proteins undergo when they fold from a disordered state to a specific three-dimensional structure reflect the evolutionary history of proteins. When we engage in reverse thinking, it will be a feasible way to exploit protein folding kinetic information and predict the protein folding pathway by exploring the evolutionary history of proteins through multiple structures alignments of family proteins. Moreover, after AlphaFold2 and ESMFold made breakthroughs in the protein structure prediction, DeepMine and Meta teams released structure databases of 214 million and 617 million, respectively^28, 29^. The availability of large structure databases can undoubtedly provide valuable data for the prediction of protein folding pathway.

In this work, we developed FoldPAthreader, an *in-silico* method for predicting protein folding pathway. This work builds on PAthreader advances by exploiting folding information from 100-million-level structure databases to design folding force field model for guiding protein folding simulations^30^. The method not only identified the folding intermediates free of any arbitrary thresholds or parameters, but also successfully predicted a series of transition states from the amino acid chain to the native state. We quantified the results using the lDDT evaluation metric^31^. The results reveal the close link between protein evolution history and folding mechanism. This work demonstrates that FoldPAthreader has developed into an effective tool for quantitative computational studies of protein folding and dynamics, which can provide a complement to experimental techniques. To the best of our knowledge, this work is the first folding force field model developed specifically for protein folding pathway prediction. It comprehensively uses the state-of-the-art modeling method AlphaFold2^28^, the fastest structure search tool Foldseek^32^ and the most abundant structure database AlphaFold DB^33^.

## Results and Discussion

### FoldPAthreader overview

The pipeline of FoldPAthreader is shown in **Fig. 1**, and the details are described in Methods. Starting from the query sequence of the target protein, the three-dimensional structure is first modeled by AlphaFold2, and remote homologues of the target are searched from the AlphaFold DB50 library through the fast structure search method Foldseek^32^. Then, structures with TM-score ≥ 0.3 are selected for multiple structures alignment (MSTA). Based on different distance deviation thresholds, the residue frequency score (ResFscore) is calculated from the MSTA, which is further designed as a statistical potential energy function combined with the residue distance information extracted from the predicted structure. Meanwhile, a folding fragment library, completely different from the modeling fragment library, is constructed based on the structures screened from MSTA, which implicitly contains folding information and is specifically used for folding pathway prediction. Finally, the protein folding pathway is simulated through three different stages of Monte Carlo conformational sampling based on fragment assembly guided by statistical and physical potential energy force fields with different energy terms and weights.

**Fig. 1.**
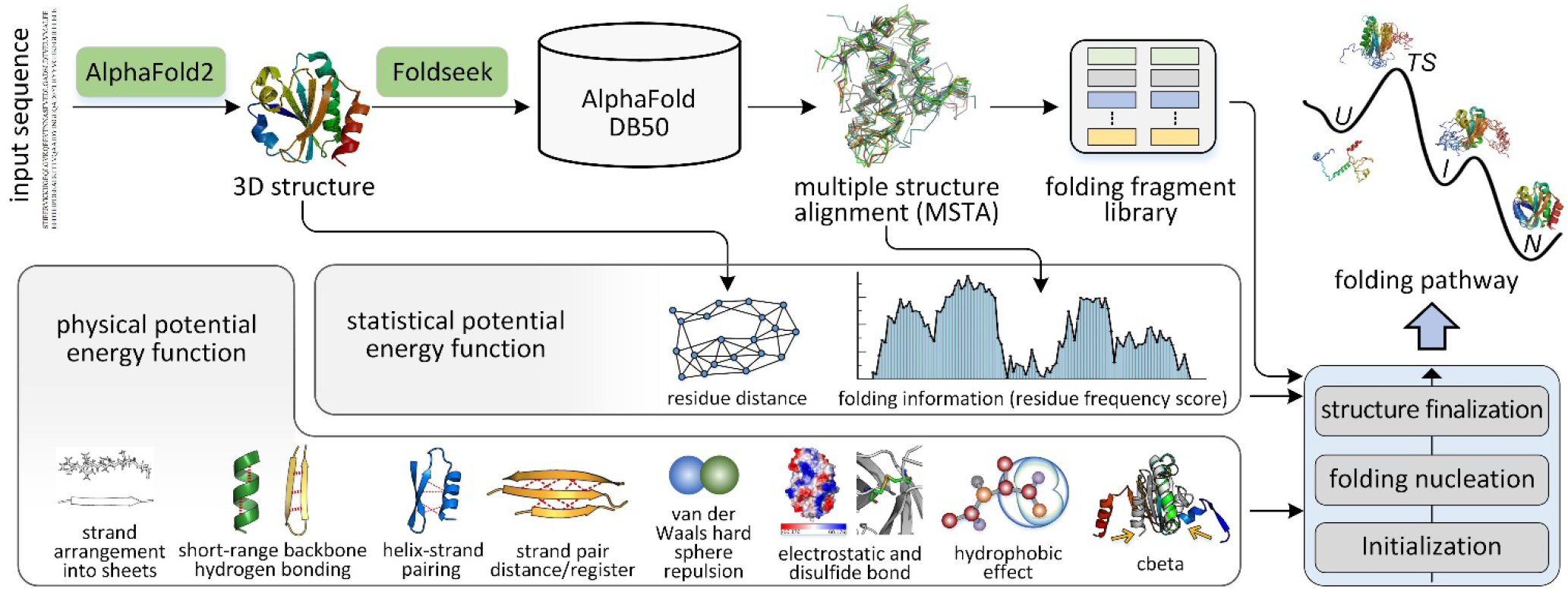
Overview of the FoldPAthreader workflow. The pipeline consists of six consecutive steps: 3D structure modeling and residue distance extraction, homologous structure search, multiple structures alignment, folding information extraction and fragment library generation, statistical and physical potential energy function construction, and folding pathway prediction.

### Comparison with biological experimental data

We collected 30 proteins to test the performance of FoldPAthreader on folding pathway prediction. Details are listed in ***SI Appendix*, Table S1**. These proteins have been analyzed by experimental techniques such as circular dichroism^34^, hydrogen deuterium exchange mass spectrometry^35^, cross-linking mass spectrometry^36^, and fluorescence resonance energy transfer^3^ to obtain relevant information that can describe the folding mechanism, including intermediates, transition states. From the literature, we found evidence and descriptions of the folding order of 30 proteins and presented them in the ***SI Appendix*, Text S1**. The experimentally determined folding order is shown in **Fig. 2** with different colors. The blue regions are first folded, followed by the gray. Experiments and molecular dynamics studies generally focus on detailed investigation of one protein at a time, with each study performed under different conditions or using different techniques^12^. We performed multiple analyzes on this dataset that focused on elucidating basic principles of protein folding without discussing the physicochemical properties of each individual protein in detail.

**Fig. 2.**
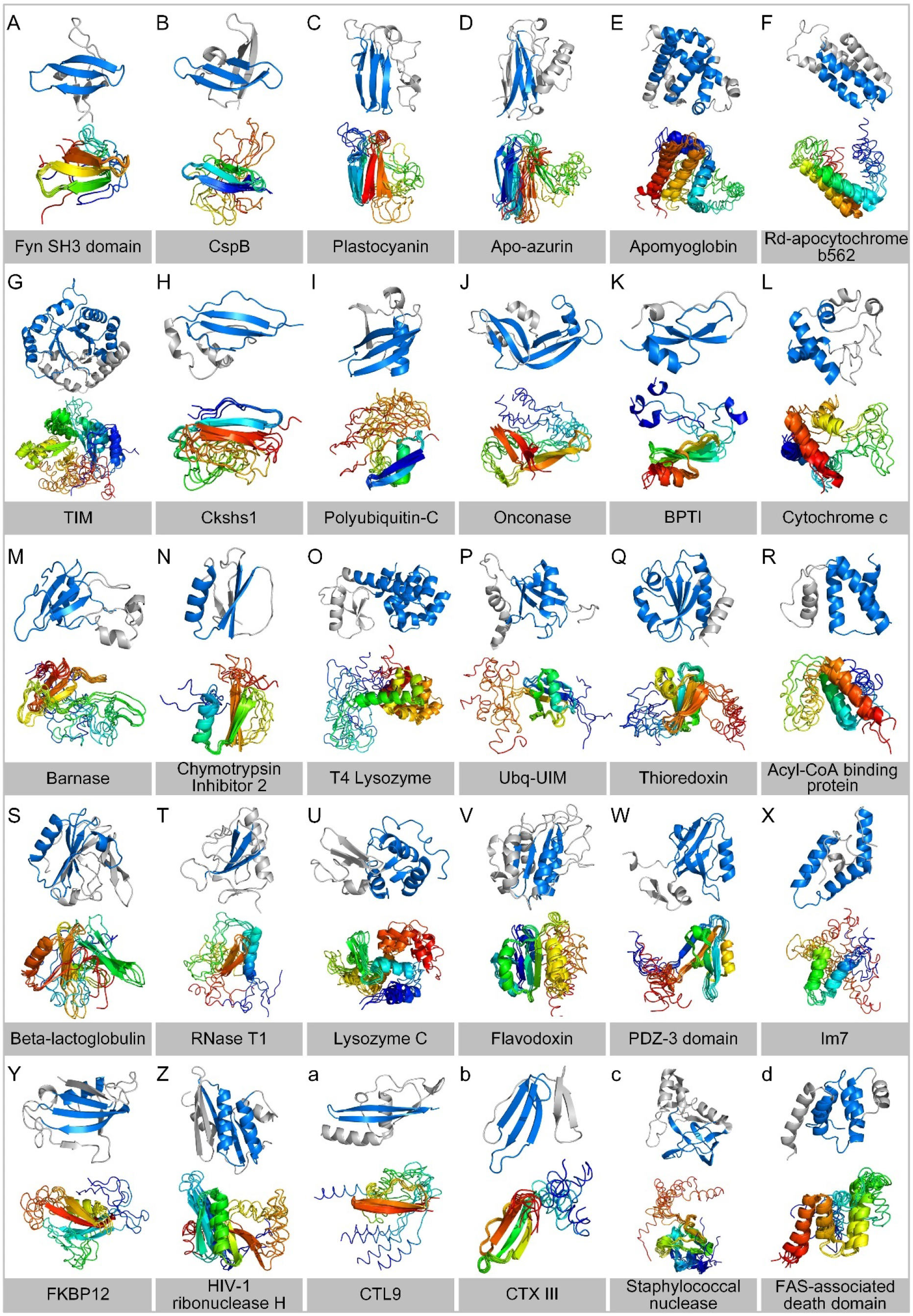
The results of 30 proteins. The blue-grey structure is an annotated folding order in the native state based on the description in collected reports. The blue regions are first folded, followed by the gray. The colored structures are the intermediates ensemble simulated by FoldPAthreader. Their partial structures with high overlap are marked with β-sheet or α-helix, indicating preferential formation.

The folding process of FoldPAthreader is divided into three stages: initialization, folding nucleation, and structure finalization. The initialization stage uses only physical potential energy functions to guide the assembly of 3-residue fragments to initialize protein chains. The simulation of folding nucleation and structure finalization stage are performed under the guidance of statistical potential and physical potential, with different weighted energy terms and number of iterations, respectively. In the folding nucleation stage, residues with higher residue frequency score form earlier constraints with other residues. Thus, the conformations of the folding nucleation stage have a tendency that the residue pairs with earlier constraints are preferentially formed. In the structure finalization stage, the weight of the energy term of ResFscore is reduced, and the overall structure is driven to fold toward the native state. Representative conformations were obtained by clustering the conformations generated from the folding nucleation stage. They are structurally superimposed as folding intermediates ensemble, and the results are shown in **Fig. 2**. The complete folding pathway of the 30 cases, including potential transition states, intermediates, and final states, are shown in ***SI Appendix***, **Fig. S1-30**. It is observed that the aligned structures of the folding intermediates ensemble are consistent with preferentially folded blue structures on most proteins compared with biological experimental data (blue-grey structures). The results suggest that FoldPAthreader has the ability to predict the folding pathway for general proteins, and is not limited to the type and size of proteins.

To further objectively evaluate the consistency of protein folding order between the predicted results and biological experimental data, we compared the lDDT of the early folded region (EFR) and the late folded region (LFR). lDDT is a scoring metric used to evaluate the local distance difference of atoms in the model, with larger values indicating greater structural similarity. It can reflect the quality of local structures at the residue level and effectively evaluate the folding order by comparing local regions of predicted intermediate and native state^31^. When the lDDT of the EFR of predicted intermediate is 10% (absolute percentage) higher than that of the LFR, it means that the early folded region forms significantly more near-native contacts than the late folded region, indicating that the folding order is consistent with biological experimental data. As shown in **Fig. 3A-B**, the blue triple-stranded β-sheet of CTX III is first folded^37^, followed by the gray double-stranded β-sheet. The lDDT of the EFR is 0.703, which is 28.4% higher than that of the LFR (0.419), indicating that the triple-stranded β-sheet of the target is preferentially formed during the folding process.

**Fig. 3.**
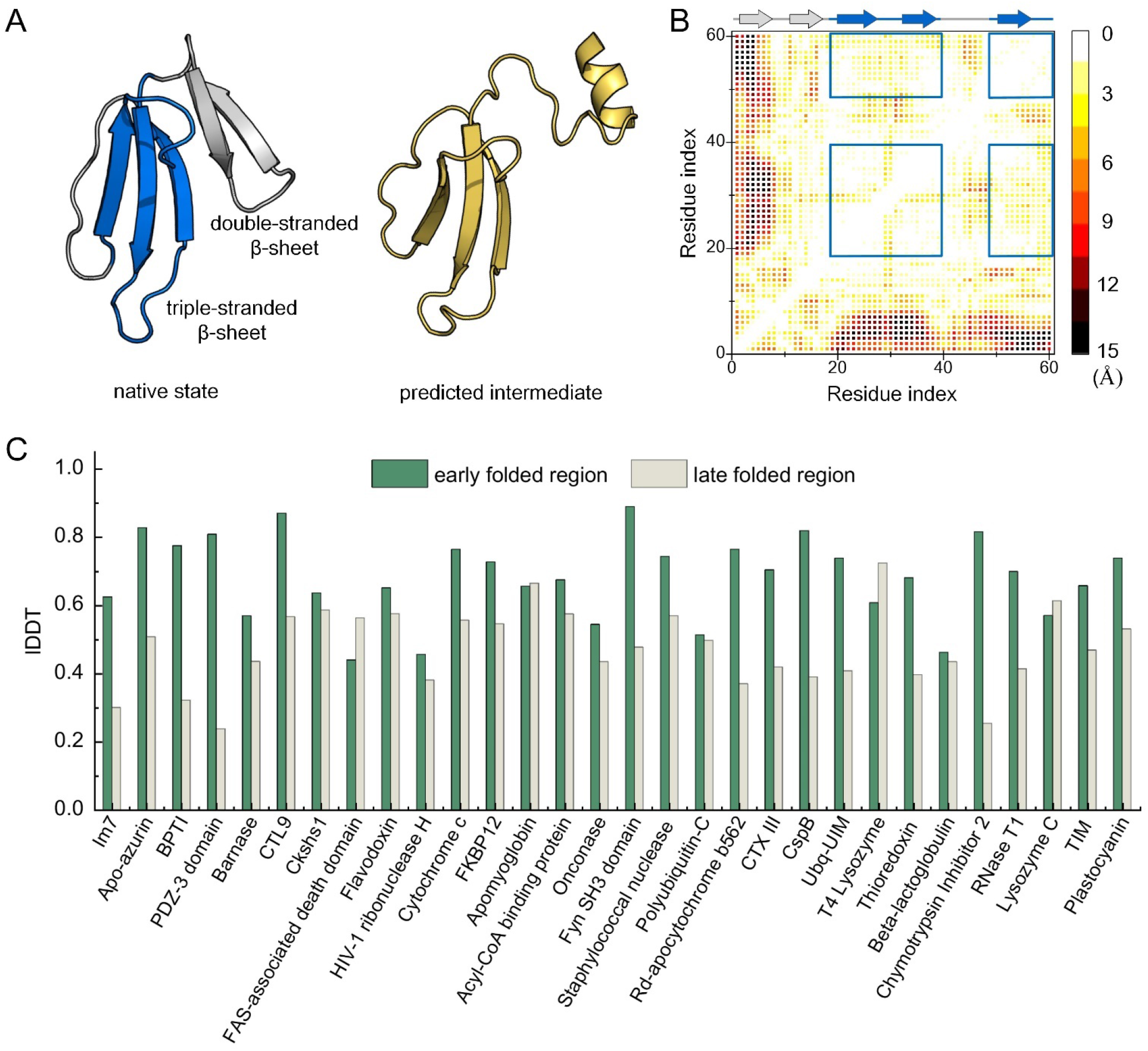
(A) The blue-gray structure is the native state of CTX III. The blue triple-stranded β-sheet are first folded, followed by the gray double-stranded β-sheet. The yellow structure is the predicted intermediate. (B) The distance difference map between the native state and predicted intermediate of CTX III. The lDDT of the EFR is calculated based on the distance difference map of the blue box, and that of the LFR is the remaining map region. (C) Comparison of lDDT between EFR and LFR of 30 proteins.

**Fig. 3C** present the predicted results of 30 cases, including 4 β-sheet proteins, 6 α-helical proteins and 20 α/β proteins. The average lDDT of EFR is 0.669 and that of LFR is 0.483. On 21 proteins, the lDDT of EFR are significantly higher than that of LFR, showing that the folding order of 70% of the proteins predicted by FoldPAthreader are consistent with the experiment data. Compared with the native state, the final state of the folding simulation has an average TM-score of 0.85, indicating that the designed folding force field model combined with the three different stage of sampling strategies are capable of folding protein to their native structure following the native folding pathway. It is interesting that the results predicted by FoldPAthreader exhibit a final state ensemble rather than a single rigid model, which may reflect the inherent flexibility and dynamism of proteins in organisms. Furthermore, MSTA-derived folding fragment libraries also contribute to accelerating the preferential formation of early folded region because the fragment libraries also contain folding information. High ResFscore regions are conserved and the derived fragments are similar, which facilitates the rapid assembly of this region. The fragments corresponding to low ResFscore regions are diverse, making the low ResFscore regions formed later in the assembly process. The predicted results support the hypothesis that protein evolutionary history implicitly contains the folding information of individual protein, illustrating that the structural space of the known protein universe is complete and can be used to study most protein folding mechanisms.

### The correlation between the evolutionary history and the folding mechanisms

Based on Ernst Haeckel’s statement^27^, we hypothesize that conformational changes in protein folding from disordered states to specific three-dimensional structures may reflect the evolutionary history of proteins. During the evolutionary, some proteins may undergo conservative changes in structure, that is, maintain similar structures during evolution because they perform similar functions. Other proteins may undergo innovative changes in structure, meaning they undergo structural remodeling to adapt to new environments or perform different functions^25^. If different species have proteins with similar structures or functions, it is often interpreted that these proteins may have evolved from a common ancestral protein^26^. Therefore, through the comparative analysis of protein structures across various species, it is possible to infer the evolutionary history of protein families, thereby enabling a deeper exploration of the folding information of individual proteins.

Here, we target plastocyanin for a detailed analysis of the correlation between protein evolutionary history and folding mechanisms. Plastocyanin is a small copper-binding protein that receive high-energy electrons from the cytochrome *b*_6_*f* complex, and then transfer these electrons to the special reaction center P700^+^ through redox reactions^38^. From a structural point of view, as shown in **Fig. 4A**, plastocyanin consists of 7 β-sheets (blue β-sandwich) and random helices (green hydrophobic patch and yellow acidic patch). Amide hydrogen exchange experiments coupled with NMR spectroscopy have demonstrated the existence of a well-populated intermediate state during the folding of plastocyanin^39^. The blue β-sheet is folded first, providing the initial context for folding. The other regions (green and yellow) then gradually converge towards the β-sandwich and form the final structure^40^.

**Fig. 4.**
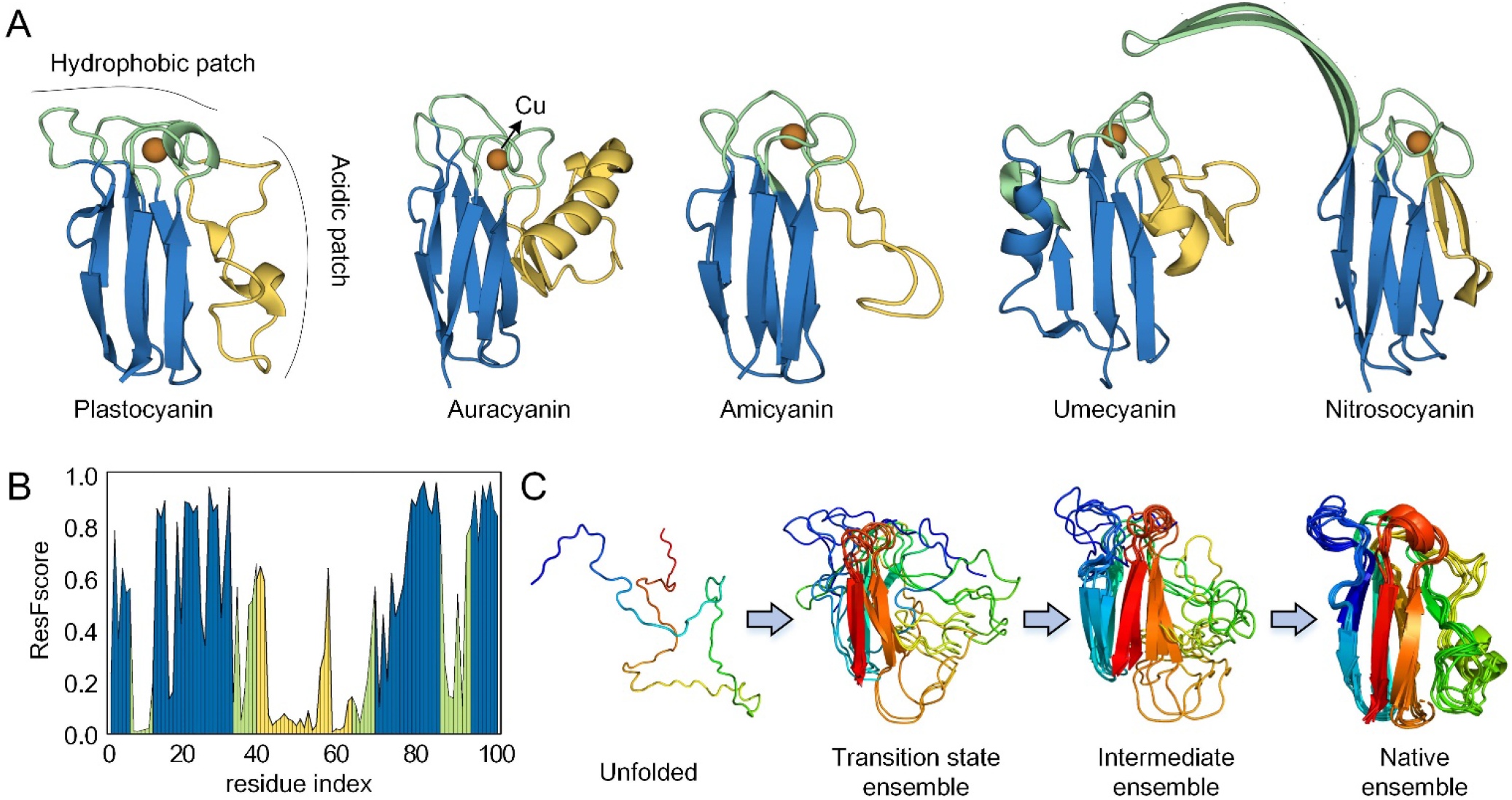
(A) 3D structure of plastocyanin (PDB ID: 9PCY) and its functionally similar auracyanin (PDB ID: 1OV8), amicyanin (PDB ID: 1ACC), umecyanin (PDB ID: 1X9R) and nitrosocyanin (PDB ID: 1IBY). (B) ResFscore distribution of plastocyanin residues obtained from MSTA. (C) The folding pathway of plastocyanin simulated by FoldPAthreader from unfolded state to native state.

From the query sequence, we predicted the folding pathway of plastocyanin protein by FoldPAthreader. In the AlphaFold DB50 structure databases, a total of 8469 structures (TM-score ≥ 0.3) were searched for globally alignment with the target protein. Then, the ResFscore of each residue is calculated, where a larger value indicates a higher frequency of residue alignment at the corresponding position of the target protein, as shown in **Fig. 4B**. It is obvious that the ResFscore of the blue region is significantly higher than that of the green and yellow regions, indicating that blue β-sandwich are present repeatedly in biological structures and are highly conserved in evolution. During the folding optimization of plastocyanin, FoldPAthreader generated a total of 709 conformations, which included gradually folded transition states and intermediates as shown in **Fig. 4C**. The transition states and intermediates ensemble show that the β-sandwich is preferentially folded, and the other regions then gradually interacts with the β-sandwich to form the final state. The results show that the folding pathway simulated by FoldPAthreader is consistent with the biological experiment^39^. This excellent performance mainly benefits from two aspects. On the one hand, the proposed folding force field focuses on the folding process, which is completely different from the traditional modeling force field such as I-TASSER and Rosetta. On the other hand, the folding fragment library can capture protein folding information from MSTA, thereby accelerating the formation of conserved regions.

In MSTA, we selected four biologically significant proteins for further analysis. These proteins are members of the Copper-bind protein family or the homologous superfamily related to Copper-bind (**Fig. 4A**). They are auracyanin from Chloroflexus aurantiacus^40^, amicyanin from Paracoccus versutus^41^, umecyanin from the roots of Armoracia rusticana^42^ and nitrosocyanin from Nitrosomonas europaea^43^, respectively. These proteins exhibit a β-sandwich architecture akin to that of plastocyanin, with differences in the Cu-binding site region and the prominent flap on the right, which is composed of helices, random loops, or β-sheets. When the four proteins were superimposed with plastocyanin, the average TM-score was 57% for the blue region but only 38% and 34% for the green and yellow regions, respectively. The similarities and differences between these structures are determined by their respective functions and processes of evolution. It has been experimentally demonstrated that the hydrophobic patch undergoes slight conformational changes when copper is removed or mercury replaces copper in plastocyanin. These conformational differences suggest a flexible region around the copper site that allows copper to be added to the folded apoenzyme^44^. As for the acidic patch, related studies have shown that it is involved in the interaction with cytochrome and contributes to rapid electron transfer in the transient complex^45^, suggesting that the hydrophobic and the acidic patch of plastocyanin are functional regions with flexibility. In the evolution of billions of years, the functional regions of proteins have undergone structural changes in order to adapt to new environmental requirements, thus deriving many homologous or remote homologous structures. For example, the nitrosocyanin monomer is part of a trimer. Its extended β-hairpins cap the copper sites of adjacent monomers, facilitating interactions through flexible conformational changes when docking with another protein^43^. For amicyanin and umecyanin, the yellow region on the right side is shorter than that of plastocyanin, and the current study has not found the functional significance of this flap. Mihwa Lee et al. concluded that it was unlikely to evolve into a smaller molecule, so it was gradually eliminated in evolution^40^. The diversity of protein structures observed within protein families is a result of evolutionary processes driven by functional selection, which reflect the evolutionary history of protein families to some extent. These pieces of evidence suggest that the correlation between protein evolutionary history and folding mechanism can be revealed from the known protein universe. That is, conserved evolutionary regions may be preferentially formed during the folding process, which is supported by our predicted results.

### FoldPAthreader folding force field captures key features of hydrogen bonding and hydrophobic interactions

In structural bioinformatics, protein hydrogen bonding and hydrophobic interactions have always been considered the key features for determining protein folding and stability^10^. In this work, in addition to the statistical potential energy function, hydrogen bonding and hydrophobic interactions are also included in the folding force field to capture key features of folding dynamic.

We found that the results of FoldPAthreader were different for proteins with different secondary structure types. As shown in ***SI Appendix*, Table S2**, the average lDDT of the EFR for β-sheet, α-helix and α/β type protein intermediates are 0.778, 0.654 and 0.652, respectively, which are 32.4%, 14.9% and 19% higher than the LFR respectively. It is obvious that the EFR of β-sheet folds the fastest, whereas the LFR of α-helix seems to fold faster than both the β-sheet and α/β type proteins, indicating that the folding priorities of proteins with different secondary structure types are different. One probable explanation is that hydrogen bonding interactions accelerate the formation of β-sheets and α-helices. Both the β-sheet and α-helix utilize hydrogen bonding to maintain their specific secondary structures, but the arrangement of the polypeptide chains and the locations of the hydrogen bonds are distinct between the two structures. The hydrogen bonds in β-sheet are formed between the carbonyl oxygen of one strand and the amino hydrogen of an adjacent strand, which can be either parallel or antiparallel^46^. The β-sheets or β-barrels formed by the multi-strand β are very tightly bound, and their structures are stable and evolutionarily conserved, making it highly likely that β-sheets are formed preferentially during the folding process. In the α-helix, the hydrogen bonds are formed between the carbonyl oxygen atom of one residue and the amino hydrogen atom of a residue located four positions down the chain^47^. This regular pattern of hydrogen bonds stabilizes the helical structure so that individual helices may preferentially fold. But the stable interaction between helix and helix might take more time to establish. This is also supported by the predicted result that LFR of α-helical proteins has higher IDDT than other types.

To further analyze the effect of hydrogen bonding interactions in folding, we calculated the proportion of secondary structure in the conformations during the initialization stage. The initial conformations of 30 proteins contained an average of 28% helical and 3% sheet structure, which is basically consistent with the results of 12 proteins simulated by David E. Shaw et al. using Anton^12^. They reported that the initial conformation contained 16% helical and 5% sheet structure. Although the data sets are different, the FoldPAthreader results exhibit the same tendency as the MD simulations in that the proportion of helices is higher than sheets in the early folding stage, indicating that individual α-helix are formed instantaneously and much faster than individual β-sheet in the early folding stage. These results again demonstrate that FoldPAthreader is effective as well as significantly less computationally expensive than MD simulation.

In addition, early studies have emphasized the importance of distinguishing between solvent-exposed and non-solvent-exposed residues in understanding protein structure and function^48^. Here, we investigated the effect of hydrophobic interactions on protein folding nucleation by calculating the relative solvent accessibility (RSA) of residues in EFR and LFR using DSSP^49^. The RSA value of a residue is obtained by dividing the absolute accessible surface area by the residue-specific maximum accessibility value^50^. If the RSA was below 25%, the residue was classified as buried residue, otherwise it was classified as exposed residue. The results are shown in ***SI Appendix*, Table S3**. The buried residues of EFR and LFR in the native structure are 53.2% and 39.6%, respectively, and the that of intermediate predicted by FoldPAthreader are 38.4% and 26.7%. The buried residues of the intermediates by FoldPAthreader are lower than the biological experimental data, which can be explained by the fact that the intermediates are not fully folded, resulting in more residues being exposed in solution. However, both sets of data show that EFR have higher buried residues than LFR, suggesting that hydrophobic amino acids are more prevalent in the EFR. This is consistent with experimental reports that proteins typically form a hydrophobic core region during folding^51^, which reduces the free energy of the system and thus promotes further folding of the protein toward its native state. Overall, the results indicated that the folding force field of FoldPAthreader can capture key features of protein folding dynamics such as hydrogen bond and hydrophobic interactions, demonstrating FoldPAthreader’s ability to predict folding pathways.

### The folding process is conserved in homologous proteins

Some studies have reported that protein folding rates are dependent on native topology, that is, proteins with similar structures often have same folding rates even if the sequences are different^21, 52^. This suggests that the folding mechanism may be conserved among homologues, meaning that they may have similar intermediate states or transition states during protein folding. In the datasets we collected, CspB and Fyn SH3 domain have been reported to be homologues^53, 54^. CspB is a 67-amino acid cold-shock protein from Bacillus subtilis that helps cells survive at low temperatures^55^. The Fyn SH3 domain is a protein domain consisting of 59 residues, which exists in a large number of eukaryotic proteins involved in signal transduction and cell polarization^56^. The sequence identity of the two proteins is only 22.4%, but they are similar in structure. As shown in **Fig. 2A** and **B**, they are composed of five β-strands arranged as two tightly packed antiparallel β-sheets, forming a closed β-barrel structure. The difference is that the triple-stranded β-sheet of CspB is composed of β1-β3, while that of Fyn SH3 domain is β2-β4. There is already sufficient evidence in the existing literature that the folding mechanisms between CspB and Fyn SH3 domain are similar, with folding intermediates characterized by folded triple-stranded β-sheet and unfolded remaining regions^21^. In the predicted results, the intermediates ensemble of CspB and Fyn SH3 domain are both well aligned on the triple-stranded β-sheet, which are consistent with the biological experimental data^55, 56^. Furthermore, the plastocyanidin and Apo-azurin in the datasets are also homologs and have similar experimental folding orders (**Fig. 2C** and **D**), that is, the β-sandwich is the preferred folding region^39, 57^. The predicted results showed that the lDDT of the EFR of plastocyanidin and Apo-azurin were 0.739 and 0.828 respectively, which were higher than the 0.531 and 0.508 of the LFR, indicating that β-sandwich of folding intermediate predicted by FoldPAthreader are preferentially formed. In general, the predicted results reveal the general principle that folding mechanisms are conserved among homologues, demonstrating that the proposed method is able to capture the potential biological properties of protein folding to some extent.

### FoldPAthreader can successfully predicted the folding pathway of BPTI and TIM

In addition to intermediates, we examined multiple transition states predicted by FoldPAthreader on the widely studied bovine pancreatic trypsin inhibitor (BPTI) and triosephosphate isomerase (TIM) proteins, whose folding pathways have been revealed by Meng Qin et al. and Kevin T. Halloran et al. using MD simulations^58, 59^. For comparison, we present the conformational snapshots of BPTI (**Fig. 5**) and TIM (**Fig. 6**) from FoldPAthreader and MD, respectively. **Fig. 5A** shows the radius of gyration of BPTI, which gradually decreases from the initial conformation to the final state. Interestingly, similar to the MD, the conformations of FoldPAthreader also temporarily fall into local basin, i.e. conformation *d*, which indicates that BPTI has intermediates in the folding process. Furthermore, it can be seen from the folding trajectory of FoldPAthreader that the conformations a-h are almost consistent with the snapshots of the conformations sampled from the MD trajectory (**Fig. 5B**). As shown in conformations *b* and *c*, the yellow β-hairpin and C-terminal α-helix adopt a native-like structure in the early stages, and the remaining regions are disordered. Next, the C-terminal helix interacts with the β-hairpin to form a stable intermediate containing two native S-S bonds, i.e. conformations *d* and *e*. Finally, the N-terminal helix gradually converges to the C-terminus through a series of transition states to form the final state (conformations *f*-*h*). The results indicated that the folding pathway predicted by FoldPAthreader for BPTI is consistent with the maximum probability pathway of MD simulation.

**Fig. 5.**
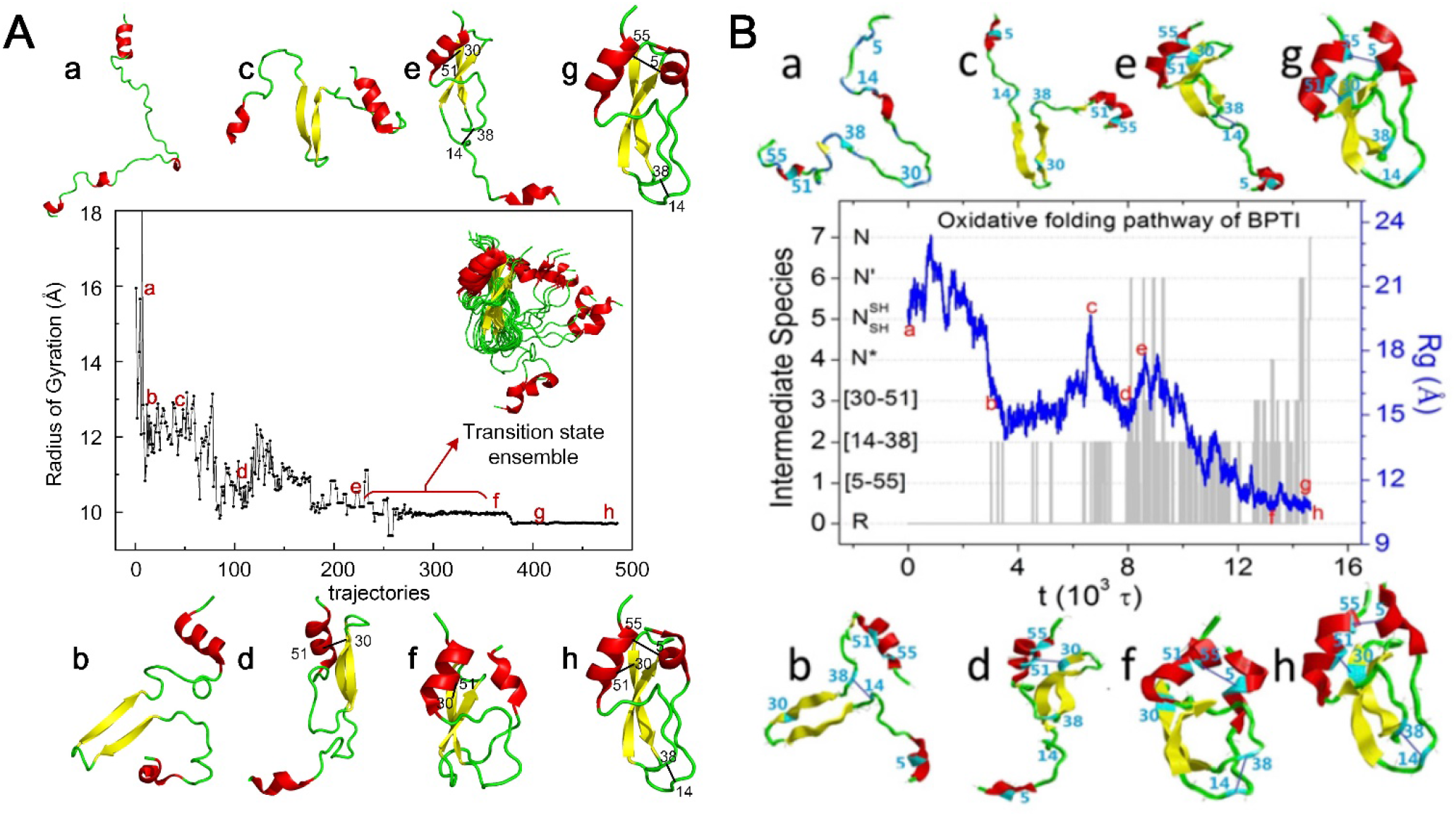
(A) 480 folding trajectories of BPTI by FoldPAthreader, showing the radius of gyration from the fully reduced starting conformation to the folded state. (a)-(h) show some of the conformations sampled in the trajectory. The right side shows the transition state ensemble from conformation (e) to conformation (f). (B) 2000 oxidative folding trajectories simulated by MD. The blue curve shows the decrease in the radius of gyration (Rg). The gray lines show formation of various disulfide species labeled on the left. Snapshots (a)-(h) show some of the conformations sampled in the trajectory. (The image B is from ref. _58_)

**Fig. 6.**
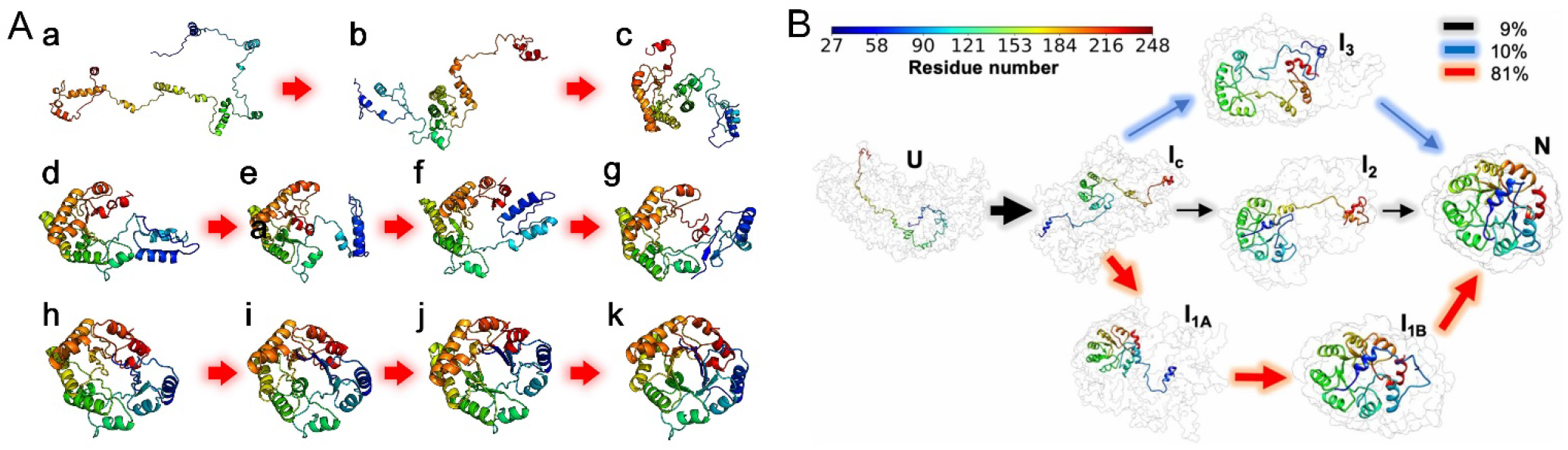
(A) Folding pathway predicted by FoldPAthreader. (B) Multiple folding pathways simulated by MD, the upper right legend shows the transition probabilities from I_c_ to I_1A_, I_2_, and I_3_. (The image B is from ref.^59^)

**Fig. 6A** and **B** present the folding pathway of TIM predicted by FoldPAthreader and simulated by MD, respectively. It can be seen that the central region of conformation *b*-*c* forms most of the contacts in the early folding stages. Then, the red region at the C-terminus forms contacts with the central region, i.e. conformation *d*-*g*. Finally, the blue region at the N-terminus converges toward the folded core to form the final state (conformation *h*-*k*). It is observed that the folding pathway predicted by FoldPAthreader is consistent with the third folding pathway simulated by MD (red arrow in **Fig. 6B**). A slight difference from the MD is the formation of the intermediates. The intermediate I_1A_ simulated by MD has a tight 7-strand barrel structure, which prevent the incorporation of N-terminus blue α and β into the barrel structure^59^. In comparison, the intermediates (conformation d-g) of FoldPAthreader exhibits a 6-strand barrel shape that includes a gap. When the N-terminus blue regions are inserted into the barrel, the overall structure becomes tighter. This suggest that there is a possibility of potential intermediates that has not been detected by MD simulation. Overall, these results again demonstrate FoldPAthreader’ ability to predict folding pathway. Compared with computationally expensive MD simulations, the FoldPAthreader protocol can be used to study the folding mechanism of large-scale proteins, greatly improving the efficiency of folding simulations.

### The performance of FoldPAthreader is related to the quality of MSTA

The excellent performance of FoldPAthreader is mainly contributed by the folding force field and the folding fragment library, which are related to the quality of MSTA. Here, we examined whether and how MSTA impact the performance of FoldPAthreader by searching for MSTA from AlphaFold DB^33^, AlphaFold DB50^32^, and Protein Data Bank (PDB) ^60^ databases respectively with the same Foldseek parameters (-s 9.5 -e 0.001 --max-seqs 10000 --alignment-type 2)^32^, and without MSTA. The AlphaFold DB database^33^, created by DeepMind and EMBL’s European Bioinformatics Institute, contains 214,683,829 entries, providing broad coverage of UniProt. AlphaFold DB50, a variant of AlphaFold DB, is a clustered database using MMseqs2 to achieve 50% sequence identity and 90% bidirectional coverage for AlphaFold DB, containing 53,665,860 structures^32, 61^. The PDB is a single global archive of three-dimensional structure data of biological macromolecules and has deposited more than 200,000 proteins as of September 2023^60^. The results of the ablation experiments are shown in **Fig. 7** and ***SI Appendix*, Table S4**. On AlphaFold DB50, the lDDT of the EFR is 0.669, which is higher than the other three performances of 0.602, 0.507 and 0.468. The number of predicted intermediates that consistent with the biological experimental data also performs best on AlphaFold DB50. This is mainly due to the fact that the homologous structures from AlphaFold DB50 are more diverse than AlphaFold DB and PDB. Although AlphaFold DB has the most homologous structures, they are extremely identical and redundant, resulting in relatively little available folding information. Likewise, the smaller PDB database structure also results in limited folding information, as evidenced by the number of effective structures (Neff-str) obtained through clustering the structure of MSTA with Foldseek and counting the number of centroids. As shown in ***SI Appendix*, Table S4**, the Neff-str of AlphaFold DB50 is 562, which is double that of AlphaFold DB and PDB, indicating that the correlation between Neff-str and precision of folding pathway is significant.

**Fig. 7.**
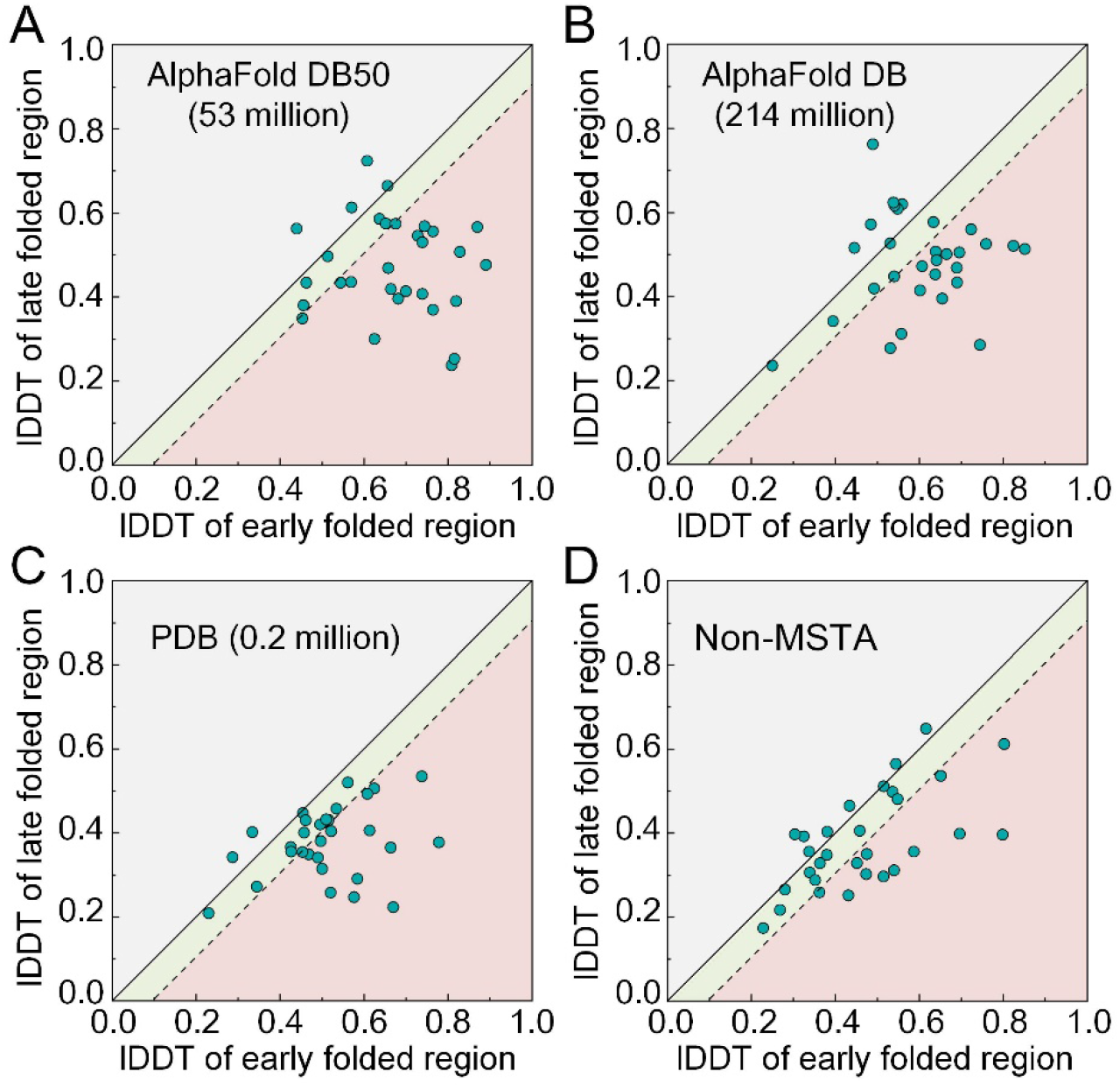
Results of MSTA ablation experiments. Head-to-head comparison between EFR and LFR of intermediates identified by FoldPAthreader using AlphaFold DB50, AlphaFold DB, PDB, and without MSTA. Red represents that the LDDT of the EFR is 10%(green) higher than that of the LFR. The cases in red areas indicate that the folding order is consistent with biological experimental data.

Furthermore, the experimental results of Non-MSTA show the folding fragment library is essential for enabling EFR to form preferentially. Non-MSTA does not use the statistical energy function from MSTA in the folding optimization, but it uses the fragment library generated by AlphaFold DB50. As shown in **Fig. 7D**, the results present that there are still 9 protein folding intermediates that are consistent with the biological experimental data even without the guidance of the statistical potential function, indicating that the folding fragment library at least partially contain folding information. In general, the diversity of MSTA determines the precision of folding force field and the quality of fragments, which together drive the protein fold to its final state following the native folding pathway.

## Discussion

At present, AI-based protein structure prediction has made a significant breakthrough. To some extent, AlphaFold2 provides only a black-box model from sequence to structure, and does not provide information about how proteins fold, which is crucial to understanding the central dogma of biology^1, 62^. In this study, we develop a protein folding pathway prediction protocol FoldPAthreader that includes a folding force field and a conformational sampling method to reveal the protein folding mechanism, which is ignored by traditional protein structure prediction methods.

This work verifies the hypothesis that evolutionary history implicitly contains the folding information of individual protein. We developed a new folding force field model and folding fragment library with folding information by searching remote homologous structures from the known protein universe (AlphaFold DB50). Different from traditional modeling, the proposed folding force field model and fragment library not only performs high-precision modeling, but also focuses on exploring folding transition states and potential intermediates. It is important to note that the final state predicted by FoldPAthreader is a conformational ensemble rather than a single model, highlighting on capturing the inherent flexibility and dynamics of the protein. The comparison with biological experimental data demonstrates that the proposed folding force field is consistent with the folding dynamics of protein in living organisms. Overall, FoldPAthreader provides a new tool for revealing protein folding mechanisms in addition to wet-lab experiments and MD simulations. The combination of physicochemical knowledge and folding evolutionary information from homologous structures will probably emerge as a new paradigm for studying protein folding mechanisms in the future.

Although proposed method achieves promising results on the given dataset, we also note some challenges. For example, proteins may contain multiple folding pathways due to the influence of environmental factors and the dynamic interactions between different regions during protein folding^3, 10^. Predicting multiple folding pathways may better capture the dynamics of the structure than a single folding pathway, but it requires the development of more complex force field models and sampling algorithms, which will be a potential direction for future research.

## Method

### Data collection

Over the past few decades, numerous wet-lab experiments have been conducted to acquire a deeper understanding of protein folding and dynamics^1^. Some progress has been made in identifying the intermediates and transition states of these proteins^63, 64^. We collected biological experimental data for a total of 30 proteins from the literature, including 4 β-sheet proteins, 6 α-helical proteins, and 20 α/β proteins, with lengths ranging from 59 to 363. Based on the collected evidence and descriptions of the folding order of these proteins, we annotated the residue range of the EFR of the protein, which has an average length of 53.7% of the total length. The residue range of some proteins may have a deviation of 1-3 residues at the boundary because some reports are not comprehensive. Detailed information is listed in ***SI Appendix*, Table S1**.

### Folding information extraction

For the input sequence, the three-dimensional structure was first predicted by AlphaFold2^28^, which was used as the input structure of Foldseek^32^ (parameters ‘-s 9.5 -e 0.001 --max-seqs 10000 --alignment-type 2’) to search for homologous structures from AlphaFold DB50^33^. The searched structures are globally aligned with the target protein through TM-align, and structures with TM-score < 0.3 are removed, which improves the quality of multiple structures alignment (MSTA). Then the frequency distribution *F* value of each residue of the target protein was calculated according to formula (1) and (2), which reflects the conservation of the protein structure during the evolution process.

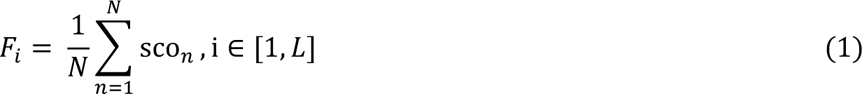

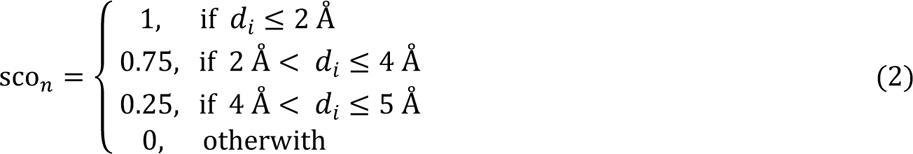

where *L* is the length of the target protein; *N* is the homologous structures number of MSTA; *d*_*i*_ is the Euclidean distance between the *i*-th residue of the target protein and the corresponding residue of the aligned MSTA structure.

### Folding force field design

The conformational sampling process of FoldPAthreader is divided into three stages, including initialization, folding nucleation and structure finalization. In the initialization stage, the physical potential energy function 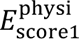 is used to guide the conformation initialization. 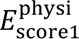 contains two energy terms: vdw and hb_srbb. The vdw term represents only steric repulsion and avoids unreasonable conformations with atomic collisions. hbond_sr_bb is the short-range backbone-backbone hydrogen bond energy term, which is to allow the helix or adjacent β hairpin to be quickly formed in the initial state. 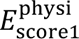 is defined as follows:

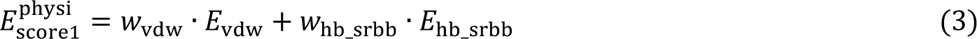

The folding nucleation stage uses physical and statistical potential energy functions. The score3 of Rosetta’s Abinitio protocol is used as a reference^65, 66^, and the pair, env, sheet, hs_pair, cbeta and rsigma terms are added to the physical potential energy function in the folding nucleation stage. 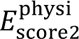 is defined as follows:

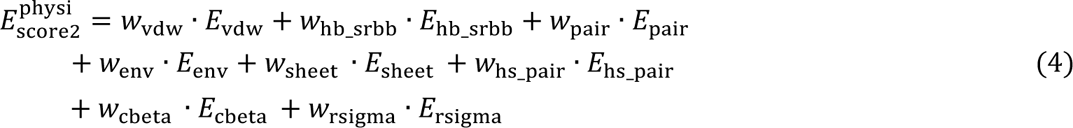

where *E*_pair_ is the energy term of the electrostatic and disulfide bond interaction of the residue pair; *E*_env_ describes the hydrophobic effect of a particular residue; the *E*_sheet_ term favors the arrangement of individual β strand into sheets. the *E*_hs_pair_ term describes the interaction between the strands and the helices. The *E*_cbeta_ is another solvation term intended to correct for excluded volume effects introduced by the simulation and favor compact structures. *E*_rsigma_ scores strand pairs based on the distance between them and the register of the two strands^67^. Different weights are used for each energy term, and the parameters are shown in ***SI Appendix*, Table S5**. The statistical potential energy function 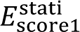 is designed based on the folding information extracted from MSTA, which is defined as follows:

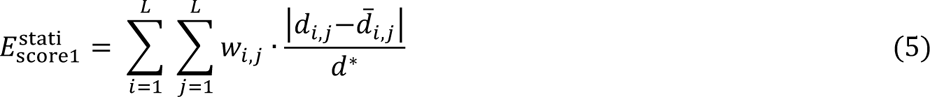

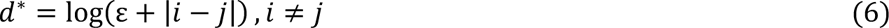

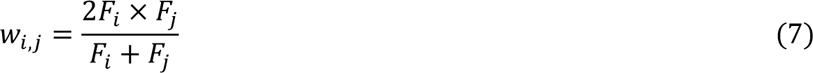

where *L* is the length of the target protein; *d*_*i*,*j*_ is the distance between the *i*-th and *j*-th residues extracted from the 3D structure of the target protein, and 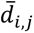 is that of the folded conformation; *d*^∗^ is the normalized scale; and ε is an infinitely small quantity so that *d*^∗^ is not zero. *w*_*i*,*j*_ is the weight for the distance deviation score between the *i*-th and *j*-th residues, which is calculated by taking the harmonic mean of *F*_*i*_ and *F*_*j*_. When both *F*_*i*_ and *F*_*j*_ are high, *w*_*i*,*j*_ will be higher. It speeds up the formation of structures corresponding to high *F* values.

In the structure finalization stage, the same physical potential energy function as in the folding nucleation stage is used, but the statistical potential energy function is different. The weight of the statistical potential energy function is removed to accelerate the region with low *F* value to converge to the folded region and form the final state. 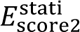 is defined as follows:

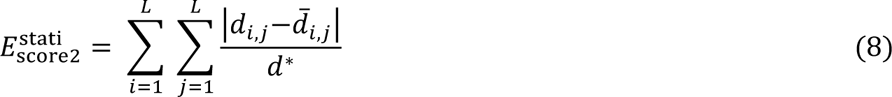

### Folding fragment library generation

The folding simulation of FoldPAthreader is based on fragment assembly for conformational sampling. Folding fragment library is a very important component for the protocol, which are derived from structures of MSTA. All structures were first ranked according to identity (TM-score) to the target protein. Then the top *M* structures are removed, and the remaining structures are used as candidate structures for generating fragments. Finally, each structure is traversed in turn, and contiguous fragments of at least 6 residues and at least 3 residues are added to the fragment list to generate a 6-residues fragment library and a 3-residues fragment library. The backbone and side chains of each fragment are represented in torsion space. *M* is defined as follows:

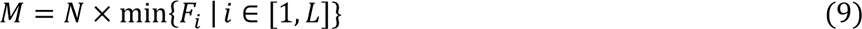

The top *M* structures have high structural overlap with the target protein, indicating that they are very identical to the target protein. The fragments generated from these highly identical structures will not carry any folding information. In contrast, the candidate structures that were screened out had locally identical or diversified regions. Candidate structure-derived fragments can avoid exploring high-energy dead ends of conserved structure regions, which accelerates the formation of conserved regions. The flexible structure regions will be assembled into more possible conformations.

### Folding optimization

FoldPAthreader uses a Monte Carlo simulated annealing search strategy for conformational sampling. In the initialization stage, the conformation is initialized by random 20\**L* times of 3-residues fragment assembly. The assembled trial conformation was scored by 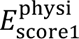, and the Metropolis criterion was used for conformational replacement.

In the folding nucleation stage, 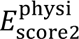 and 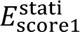 were used to score the trial conformation and the Metropolis criterion was used to select the conformation. However, the annealing temperatures of the physical potential and the statistical potential energy function are different. They are *kT*_physi_=5 and *kT*_stati_=2 respectively. The function of the physical potential energy function is to ensure that the conformation is physically reasonable, but the continuous reduction of physical energy during folding is not necessary. On the contrary, high annealing temperature can increase the probability of conformational update.

In the structure finalization stage, 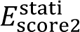 was first used to score trial conformation and Metropolis criterion was used to perform conformational replacement. If it fails, 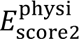 is used for scoring. This greedy search strategy speeds up the convergence of protein structures.

## Data availability

The online server and package of FoldPAthreader are made freely available at http://zhanglab-bioinf.com/PAthreader and GitHub (https://github.com/iobio-zjut/FoldPAthreader).

## Acknowledgments

This work is supported by the National Key R&D Program of China (2022ZD0115103), the National Nature Science Foundation of China (62173304), and the Key Project of Zhejiang Provincial Natural Science Foundation of China (LZ20F030002).

## Supporting Information for

### Supporting Information Table

**Table S1.** Detailed information of 30 cases. Residue ranges of early folded region were annotated according to descriptions in references. The residue range of some proteins may have a deviation of 1-3 residues at the boundary because some reports are not comprehensive.

| Protein name | PDB ID | length | Type | Residue range of early fold region | Reference (DOI) |
| --- | --- | --- | --- | --- | --- |
| BPTI | 1QLQ | 58 | $\alpha/\beta$ | 18-58 | 10.1073/pnas.1503909112 |
| Apo-azurin | 1AIZ | 129 | $\alpha/\beta$ | 27-36, 89-98, 106-112, 121-129 | 10.1002/prot.22099 |
| Im7 | 1AYI | 86 | $\alpha$ | 12-45, 66-79 | 10.1038/nsb757 |
| PDZ-3 domain | 1BE9 | 115 | $\alpha/\beta$ | 10-92 | 10.1016/j.jmb.2004.11.040 |
| Barnase | 1BGS | 110 | $\alpha/\beta$ | 1-22, 53-75, 85-110 | 10.1021/bi0362267 |
| CTL9 | 1DIV | 90 | $\alpha/\beta$ | 66-90 | 10.1002/prot.22099 |
| Ckshs1 | 1DKT | 72 | $\alpha/\beta$ | 1-21, 50-72 | 10.1016/s0022-2836(02)01202-0 |
| FAS-associated death domain | 1E3Y | 104 | $\alpha$ | 1-32, 47-79 | 10.1007/s00249-011-0756-6 |
| Flavodoxin | 1FTG | 168 | $\alpha/\beta$ | 1-8, 47-53, 80-87, 98-110, 139-143, 151-168 | 10.1006/jmbi.1998.2045 |
| HIV-1 ribonuclease H | 1HRH | 125 | $\alpha/\beta$ | 13-21, 46-72, 89-110, 125-1, 13-21, 46-72, 89-110 | 10.1002/pro.5560071014 |
| Cytochrome c | 1I5T | 104 | $\alpha$ | 1-14, 61-69, 87-104 | 10.1073/pnas.1706196114 |
| FKBP12 | 1J4H | 107 | $\alpha/\beta$ | 21-30, 56-65, 71-76, 96-107 | 10.1002/prot.22099 |
| Apomyoglobin | 1MBC | 153 | $\alpha$ | 7-18, 24-44, 61-78, 102-142 | 10.1073/pnas.0804033105 |
| Acyl-CoA binding protein | 1NTI | 86 | $\alpha$ | 1-36, 65-86 | 10.1073/pnas.0509100103 |
| Onconase | 1ONC | 103 | $\alpha/\beta$ | 17-36, 55-99 | 10.1021/bi961085c |
| Fyn SH3 domain | 1SHF | 59 | $\beta$ | 25-30, 36-41, 47-51 | 10.1073/pnas.0404436101 |
| Staphylococcal nuclease | 1STN | 136 | $\alpha/\beta$ | 1-36, 66-90 | 10.1016/j.bbrc.2008.08.073<br>10.1073/pnas.91.2.449 |
| Polyubiquitin-C | 1UBQ | 76 | $\alpha/\beta$ | 1-16, 22-34, 64-76 | 10.1073/pnas.89.6.2017<br>10.1021/bi00161a019 |
| Rd-apocytochrome b562 | 1YYJ | 104 | $\alpha$ | 22-43, 65-89 | 10.1073/pnas.0501372102 |
| CTX III | 2CRT | 60 | $\beta$ | 19-39, 49-60 | 10.1074/jbc.273.17.10181 |
| CspB | 2F52 | 67 | $\beta$ | 1-19, 46-53 | 10.1016/j.jmb.2004.04.011 |
| Ubq-UIM | 2KDI | 114 | $\alpha/\beta$ | 11-80 | 10.1016/j.bpc.2011.05.004 |
| T4 Lysozyme | 2LZM | 164 | $\alpha/\beta$ | 1-11, 65-164 | 10.1016/j.jmb.2006.10.048 |
| Thioredoxin | 2TRX | 108 | $\alpha/\beta$ | 1-90 | 10.1016/j.bbapap.2014.11.004 |
| Beta-lactoglobulin | 3BLG | 162 | $\alpha/\beta$ | 91-162 | 10.1038/84145 |
| Chymotrypsin Inhibitor 2 | 3CI2 | 64 | $\alpha/\beta$ | 12-34, 45-50 | 10.1016/j.polymer.2003.10.092 |
| RNase T1 | 3RNT | 104 | $\alpha/\beta$ | 56-61, 75-81, 86-92 | 10.1002/pro.5560060702 |
| Lysozyme C | 6LYZ | 129 | $\alpha/\beta$ | 1-39, 85-129 | 10.1110/ps.8.1.35 |
| TIM | 7TIM | 247 | $\alpha/\beta$ | 1-164 | 10.1016/s0022-2836(02)01100-2 |
| Plastocyanin | 9PCY | 99 | $\beta$ | 1-6, 13-15, 18-31, 69-83, 93-99 | 10.1021/bi00097a005 |

**Table S2.** Average lDDT of early fold regions and late fold regions of β, α and α/β type proteins.

| | | β-sheet proteins | α-helix proteins | $\alpha/\beta$ proteins | All proteins |
| --- | --- | --- | --- | --- | --- |
| IDDT | early fold region | 0.778 | 0.654 | 0.652 | 0.669 |
|  | late fold region | 0.454 | 0.505 | 0.462 | 0.483 |

**Table S3.** Proportion of buried residues in early fold regions and late fold regions. Residues with a relative solvent accessibility of less than 25% are classified as buried residues.

|  | Native of experimental |  | Intermediate of FoldPATHreader |  |
| --- | --- | --- | --- | --- |
|  | early fold region | late fold region | early fold region | late fold region |
| Buried residue | 53.2% | 39.6% | 38.4% | 26.7% |

**Table S4.** Results of MSTA ablation experiments. The number of effective structures is obtained by clustering similar structures of MSTA through Foldseek.

|  | IDDT of early folded region | IDDT of late folded region | num (consistent with experimental data) | effective structures of MSTA |
| --- | --- | --- | --- | --- |
| AlphaFold DB50 | 0.669 | 0.483 | 21/30 | 562 |
| AlphaFold DB | 0.602 | 0.585 | 17/30 | 202 |
| PDB | 0.507 | 0.373 | 15/30 | 291 |
| Non-MSTA | 0.468 | 0.382 | 10/30 | 0 |

**Table S5.** Weights of **e**nergy term for Monte Carlo conformational sampling.

|  |  |
| --- | --- |
| $w_{\text{vdw}}$ | 1.0 |
| $w_{\text{hb\_srbb}}$ | 0.5 |
| $w_{\text{pair}}$ | 1.0 |
| $w_{\text{env}}$ | 1.0 |
| $w_{\text{sheet}}$ | 1.0 |
| $w_{\text{hs\_pair}}$ | 1.0 |
| $w_{\text{cbeta}}$ | 0.5 |
| $w_{\text{rsigma}}$ | 1.0 |

### Supporting Information Figure

Figure S1-30 shows the representative conformational ensemble of the complete folding pathway from the unfolded state to the folded state of 30 test proteins, including potential transition states, intermediates, and final states.

**Figure S1.**
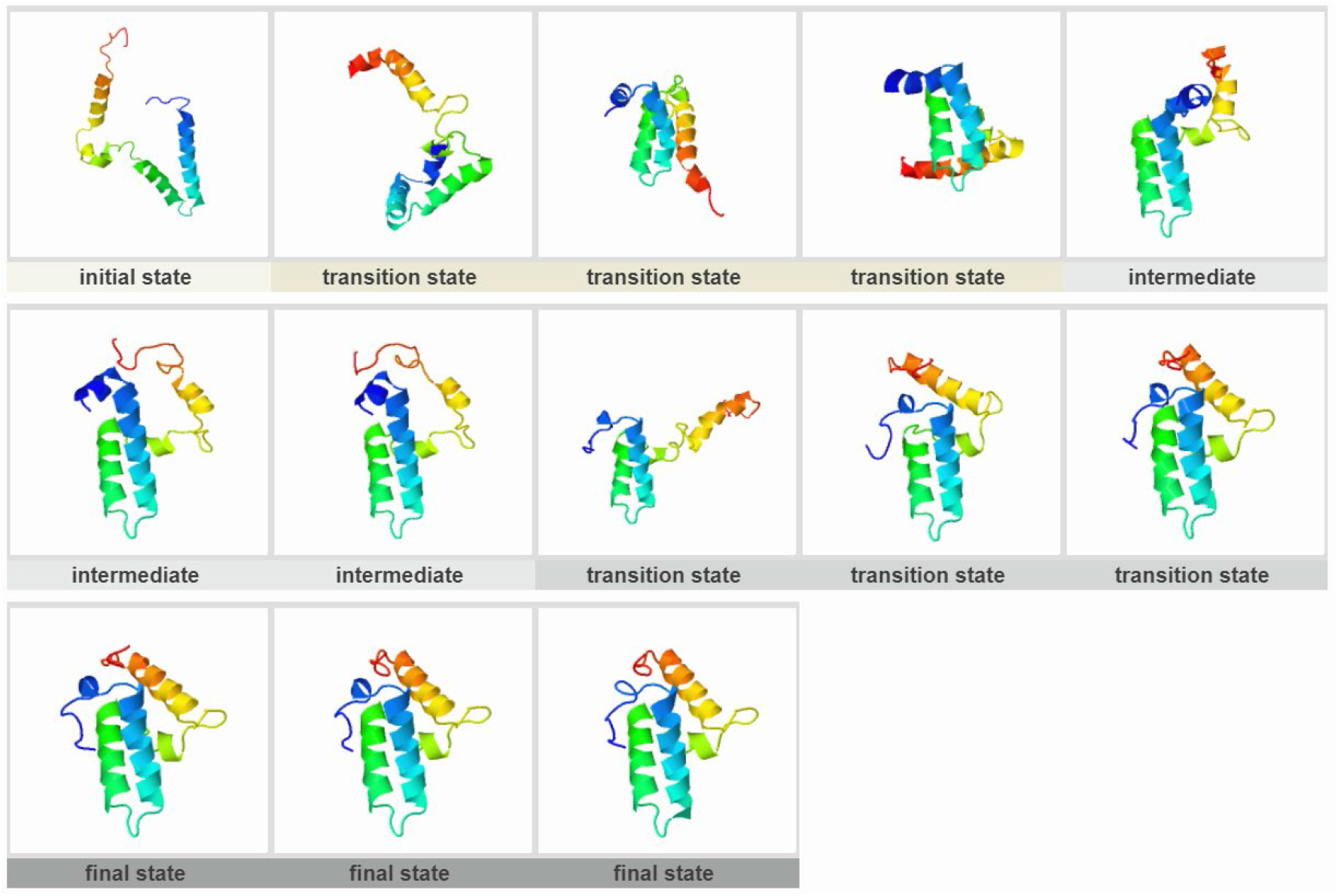
Im7 (PDB ID:1AYI)

**Figure S2.**
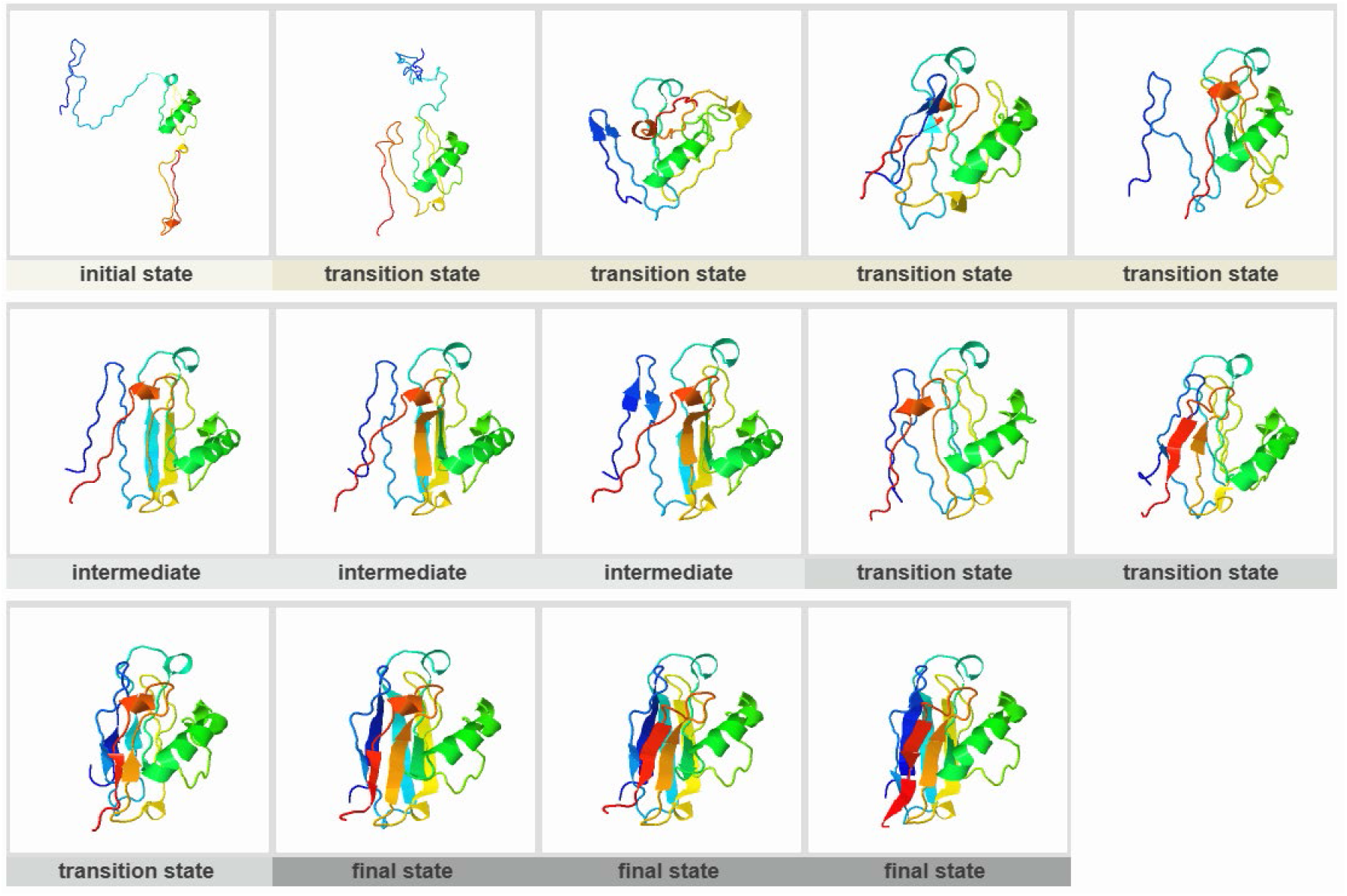
Apo-azurin (PDB ID:1AIZ)

**Figure S3.**
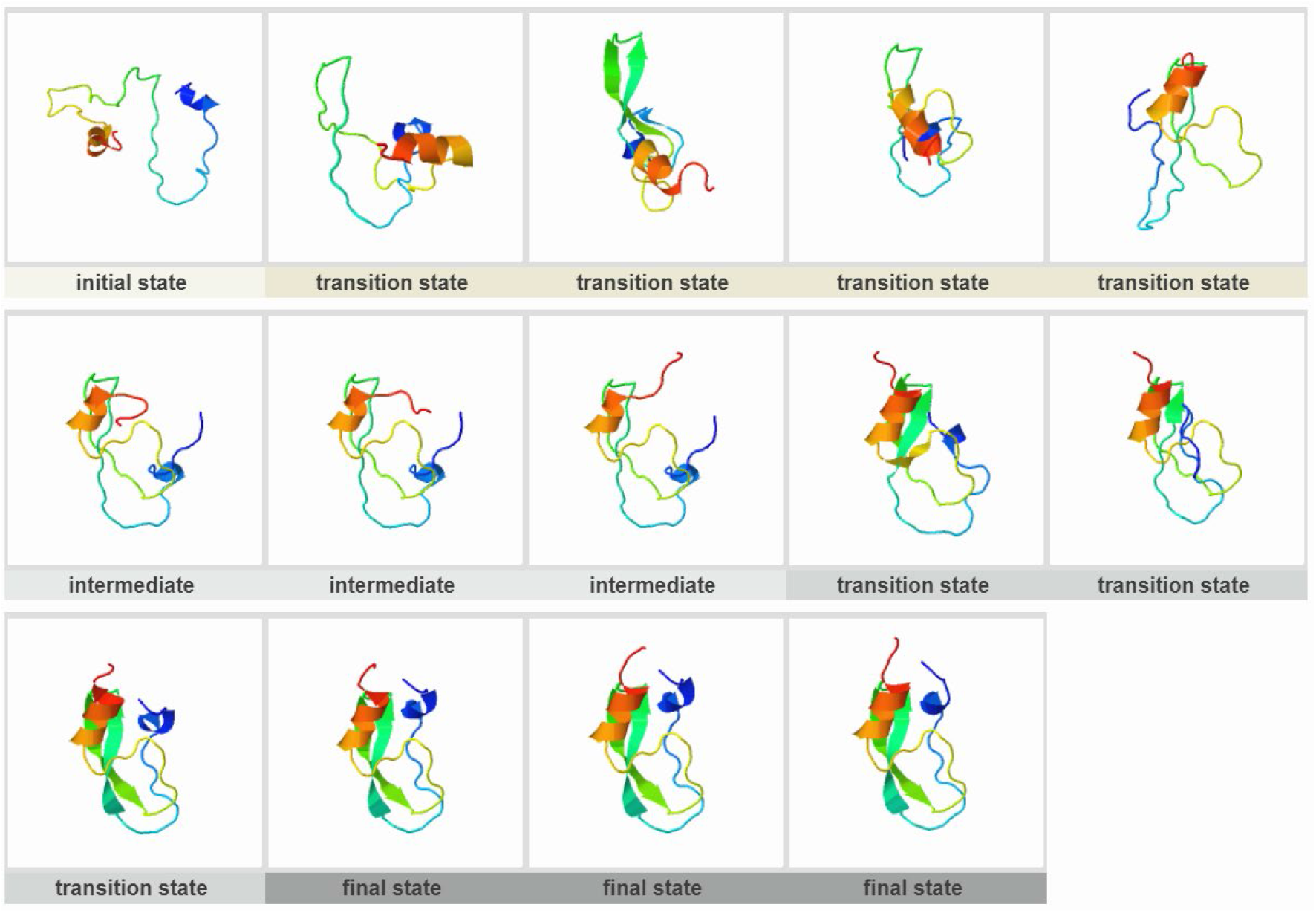
BPTI (PDB ID:1QLQ)

**Figure S4.**
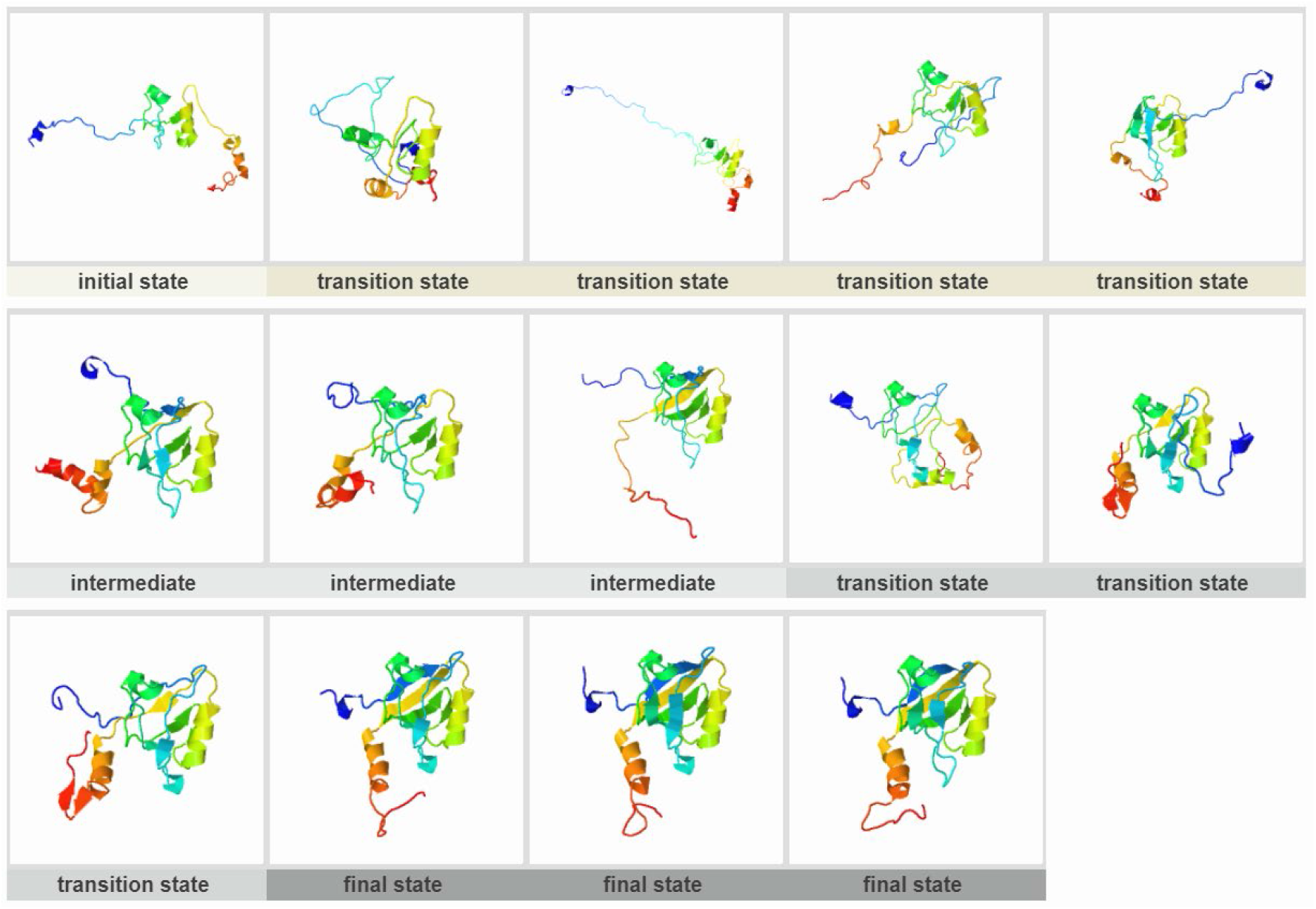
PDZ-3 domain (PDB ID:1BE9)

**Figure S5.**
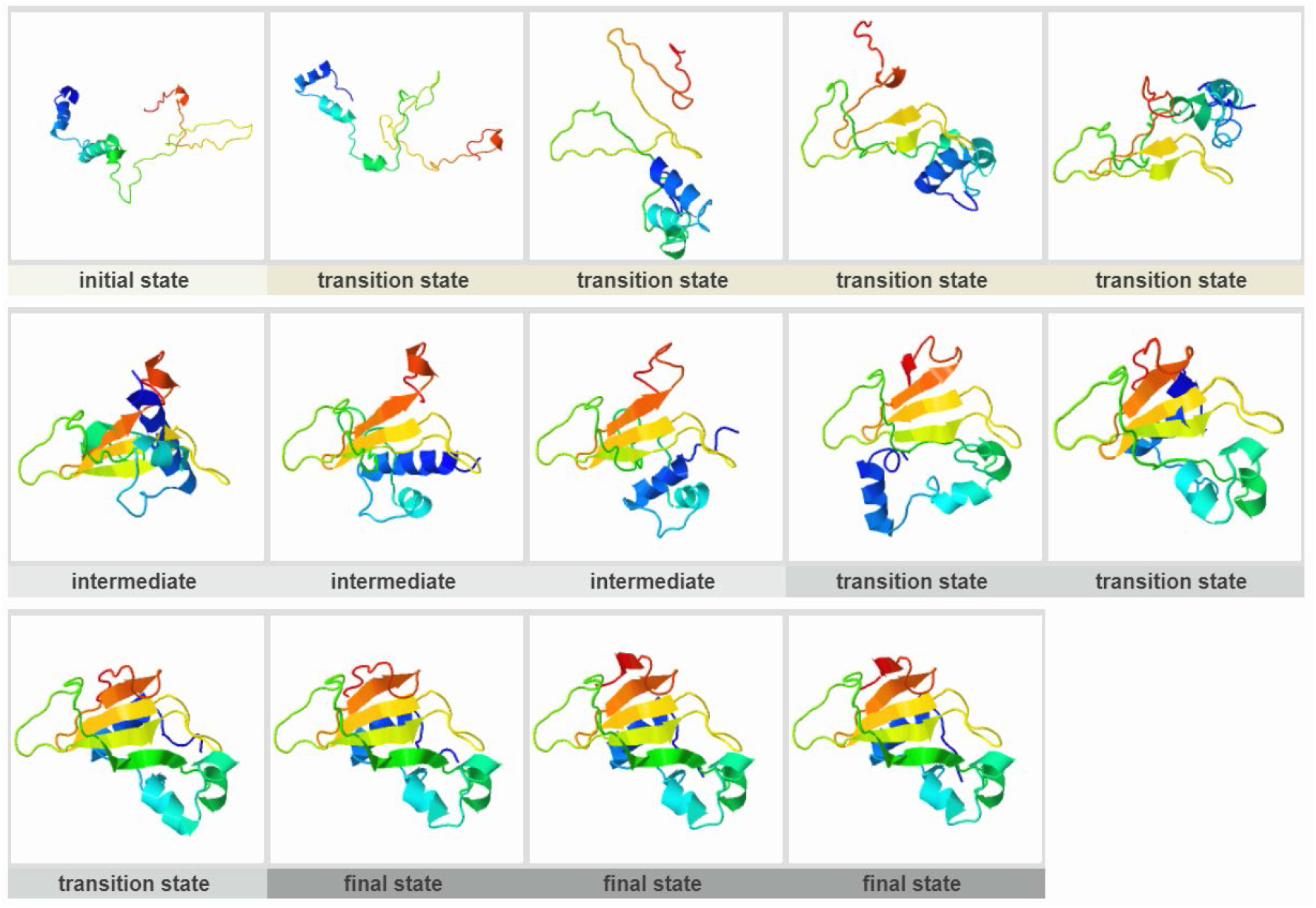
Barnase (PDB ID:1BGS)

**Figure S6.**
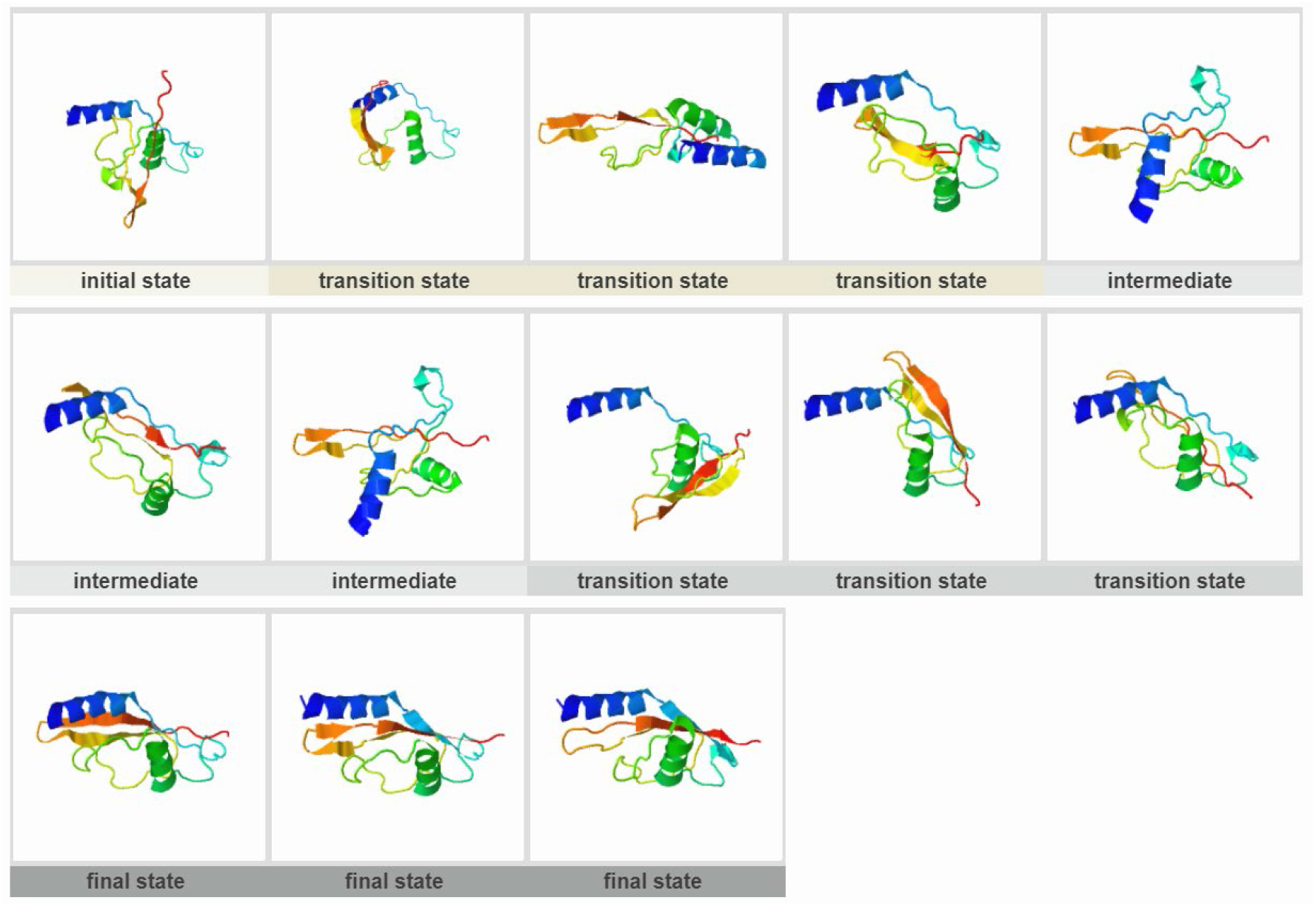
CTL9 (PDB ID:1DIV)

**Figure S7.**
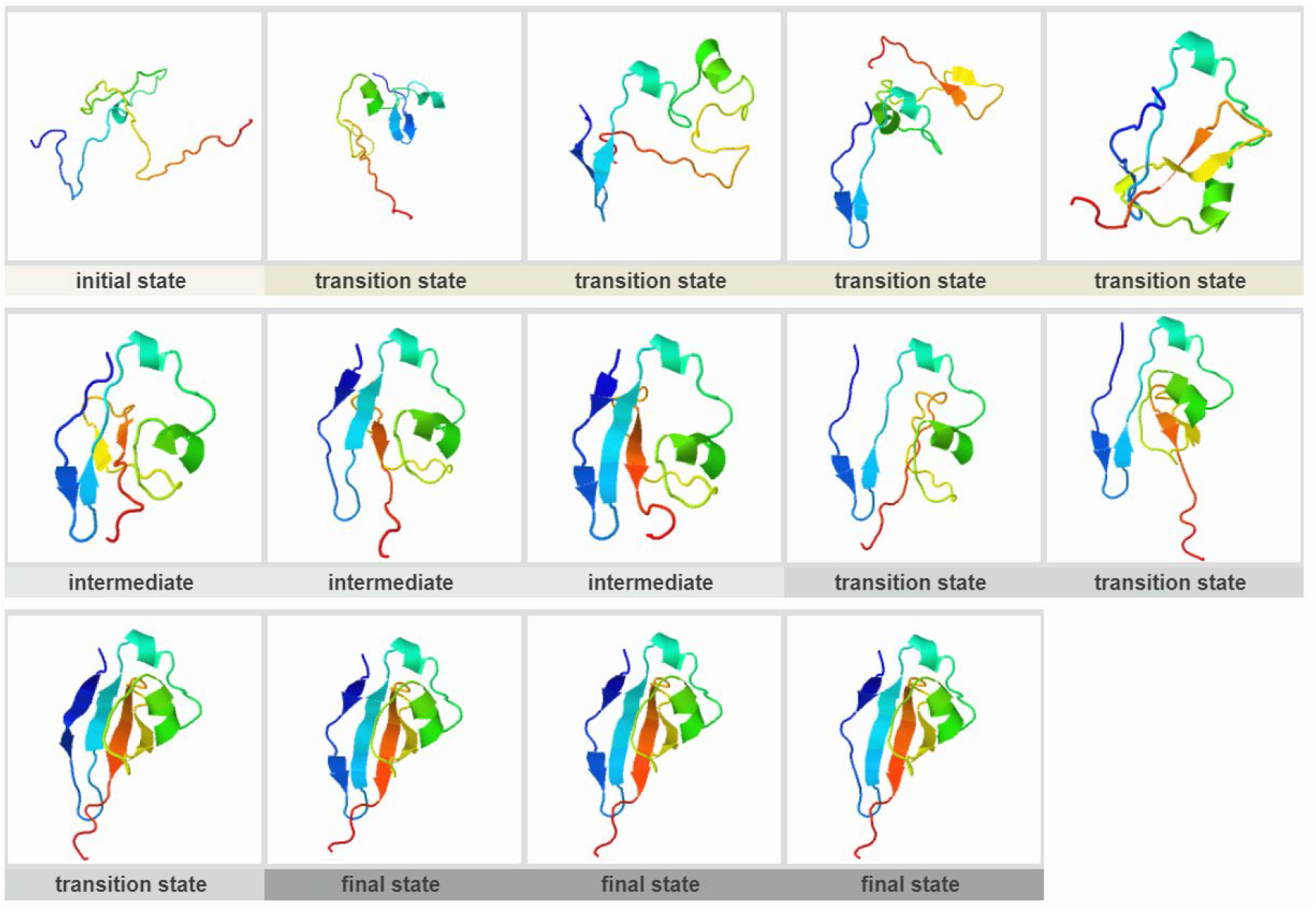
Ckshs1 (PDB ID:1DKT)

**Figure S8.**
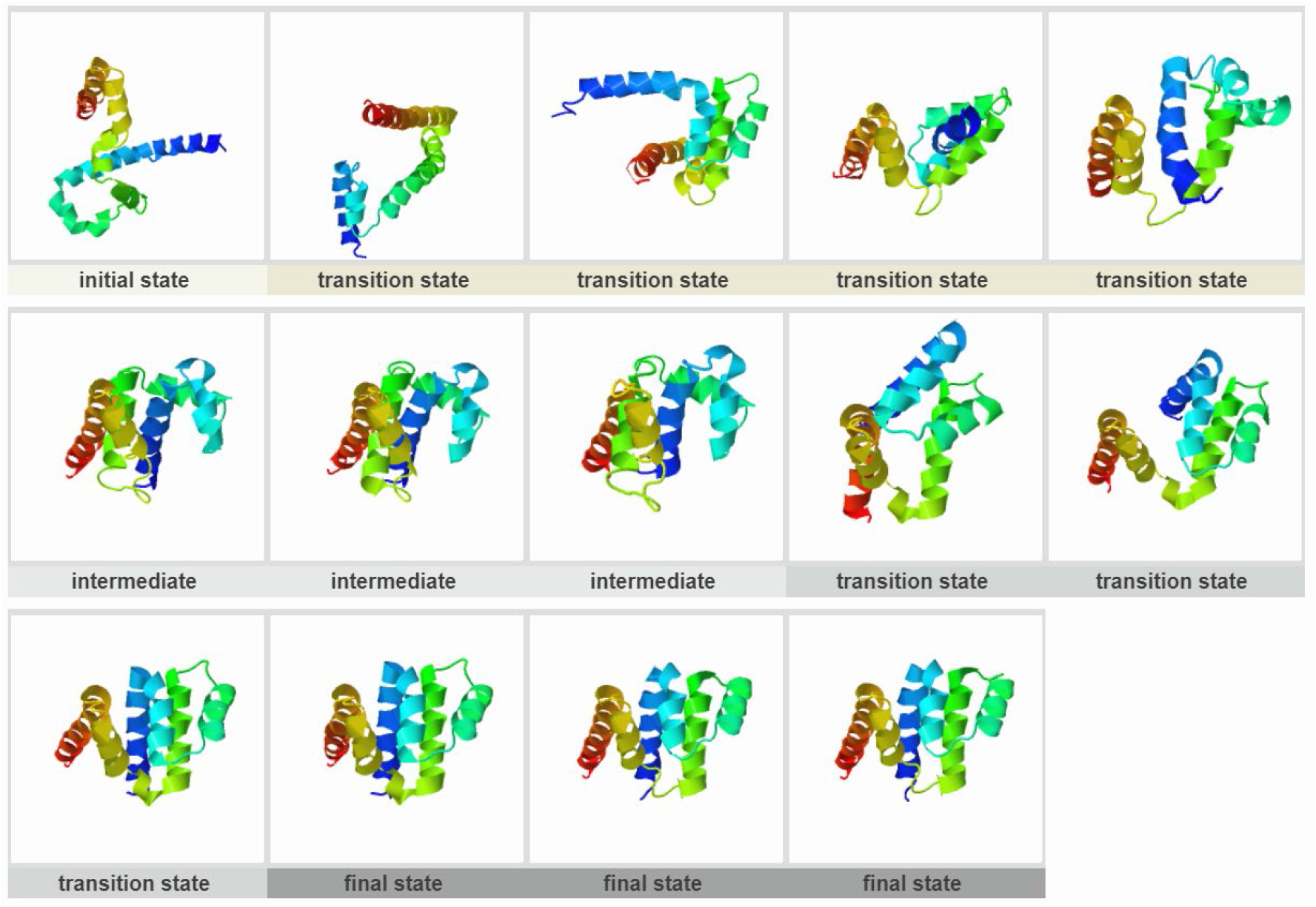
FAS-associated death domain (PDB ID:1E3Y)

**Figure S9.**
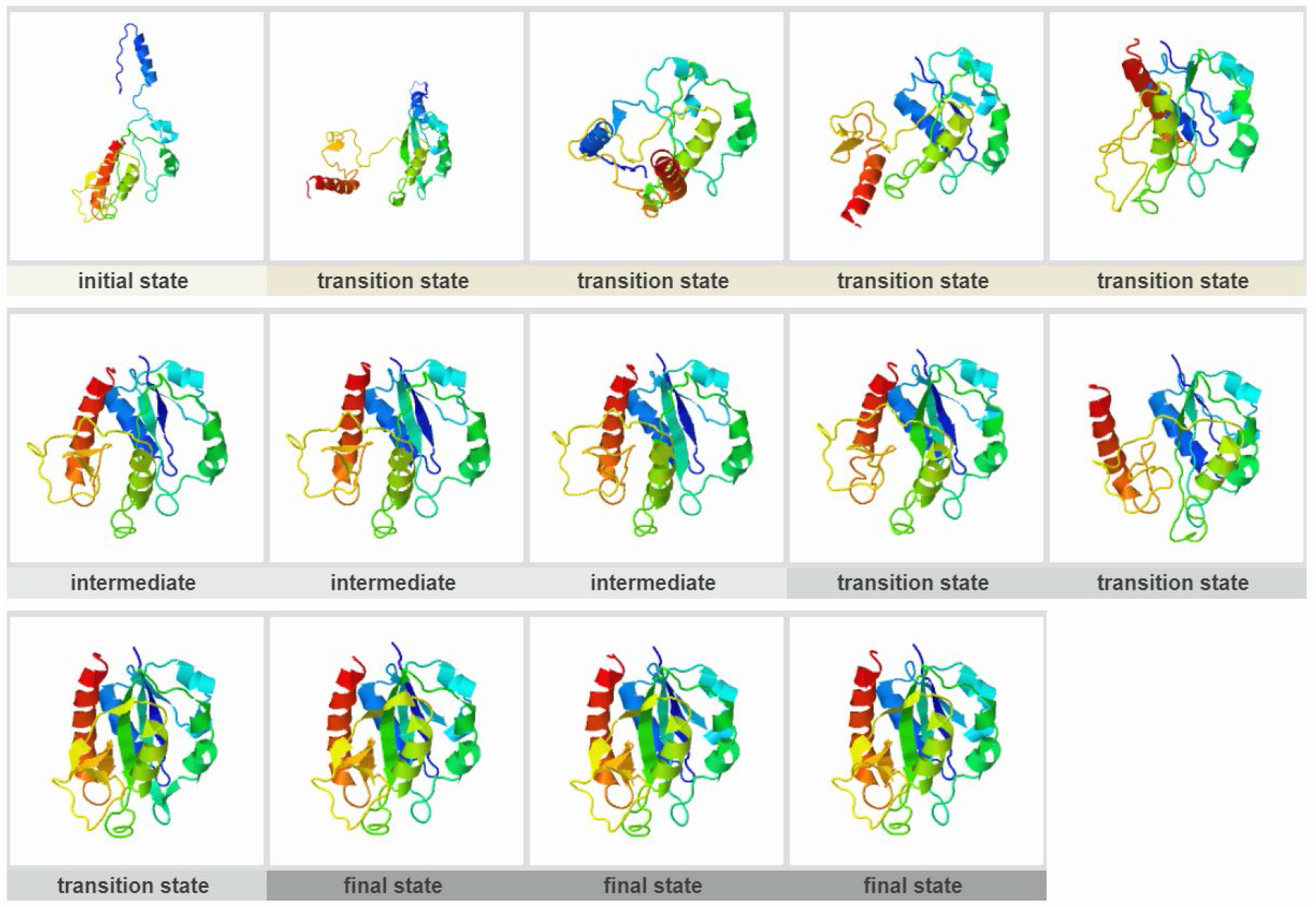
Flavodoxin (PDB ID:1FTG)

**Figure S10.**
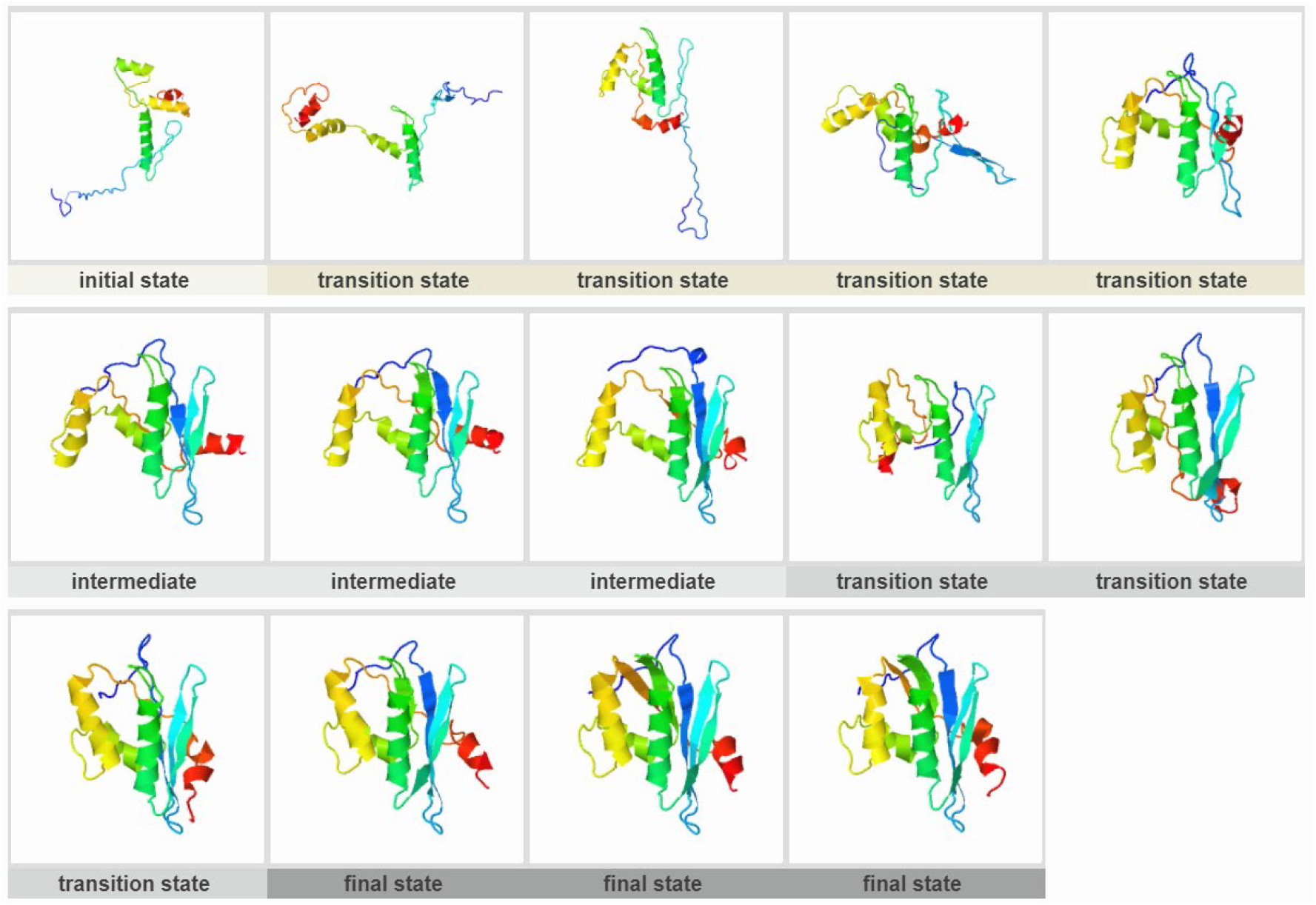
HIV-1 ribonuclease H (PDB ID:1HRH)

**Figure S11.**
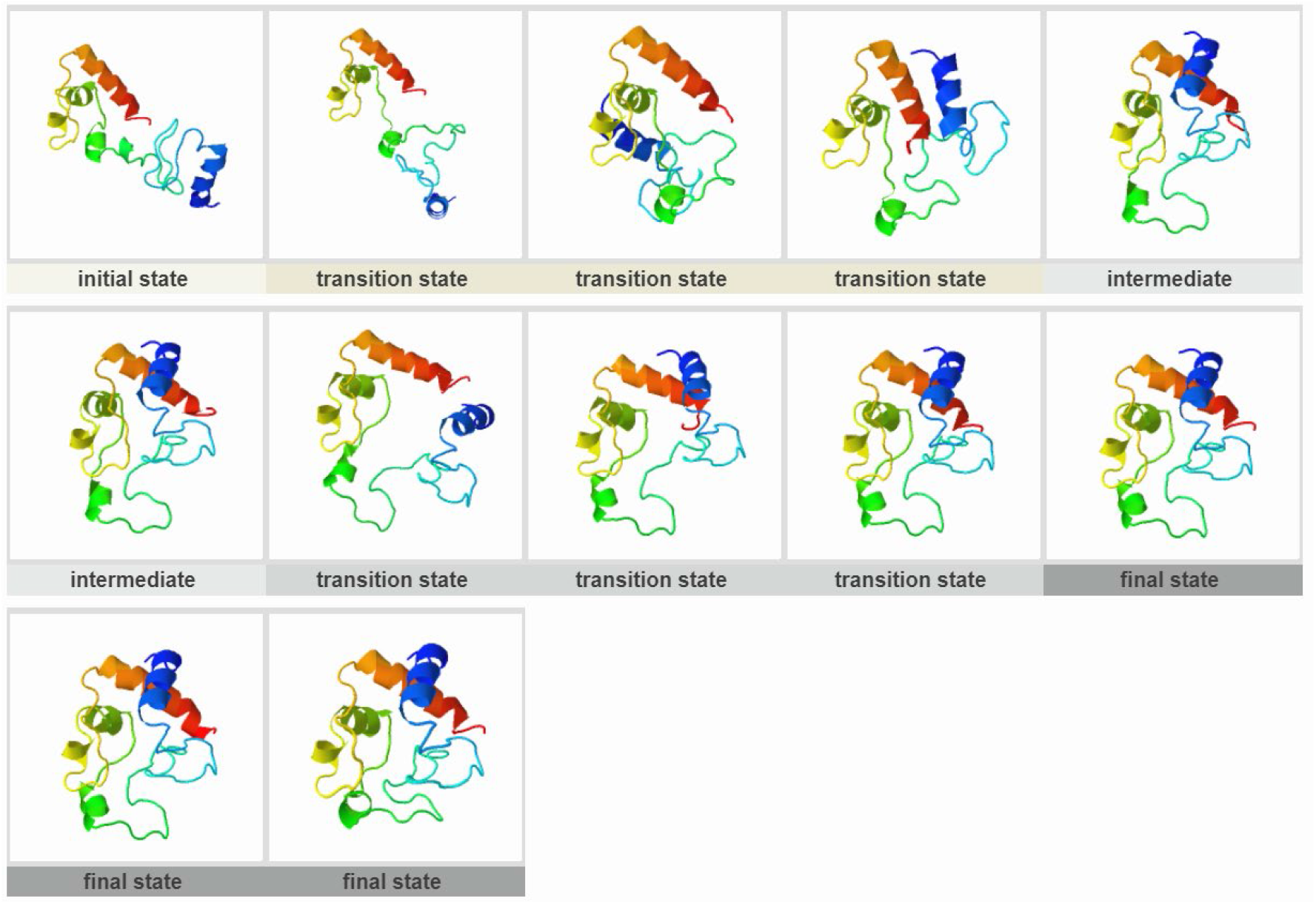
Cytochrome c (PDB ID:1I5T)

**Figure S12.**
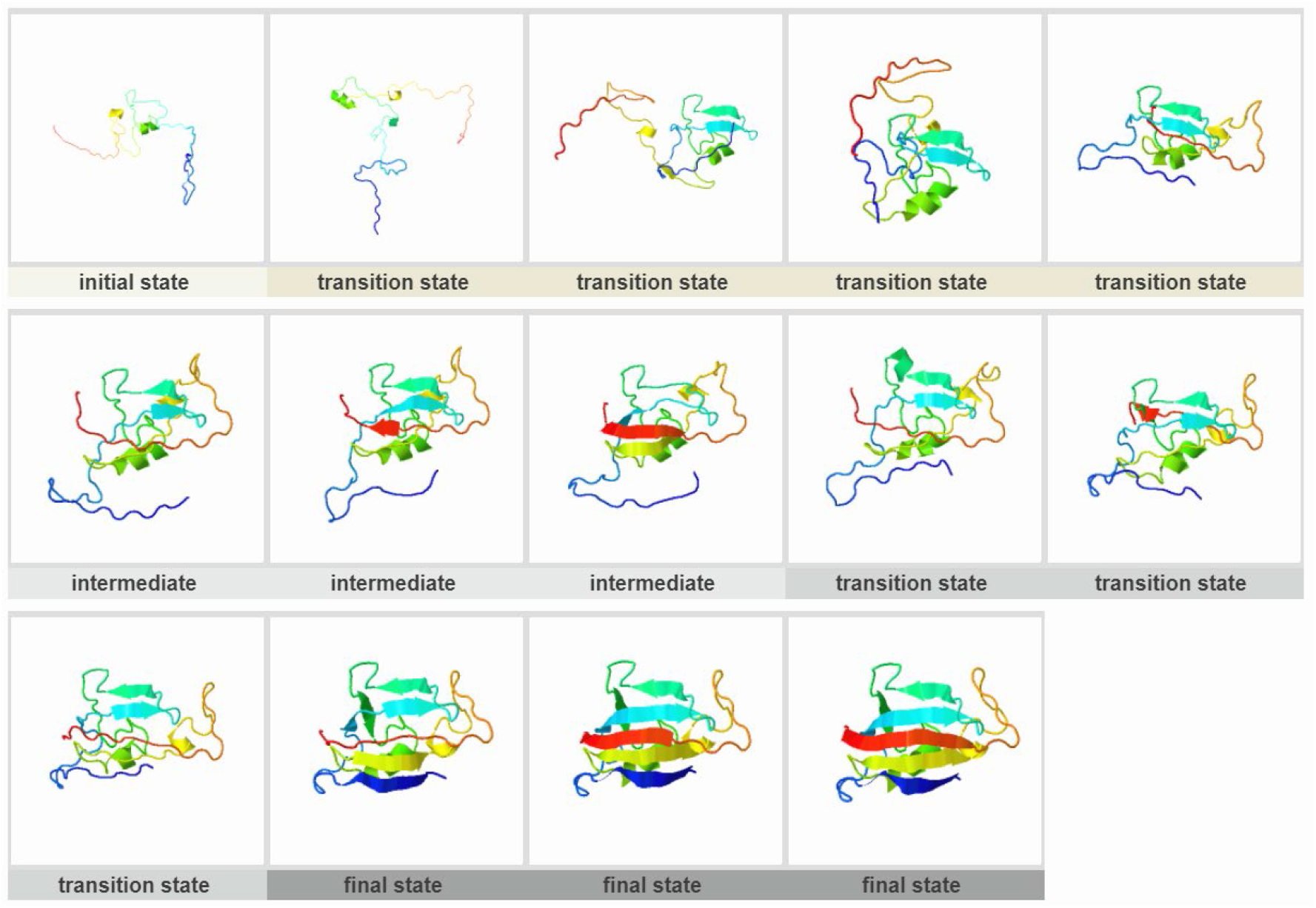
FKBP12 (PDB ID:1J4H)

**Figure S13.**
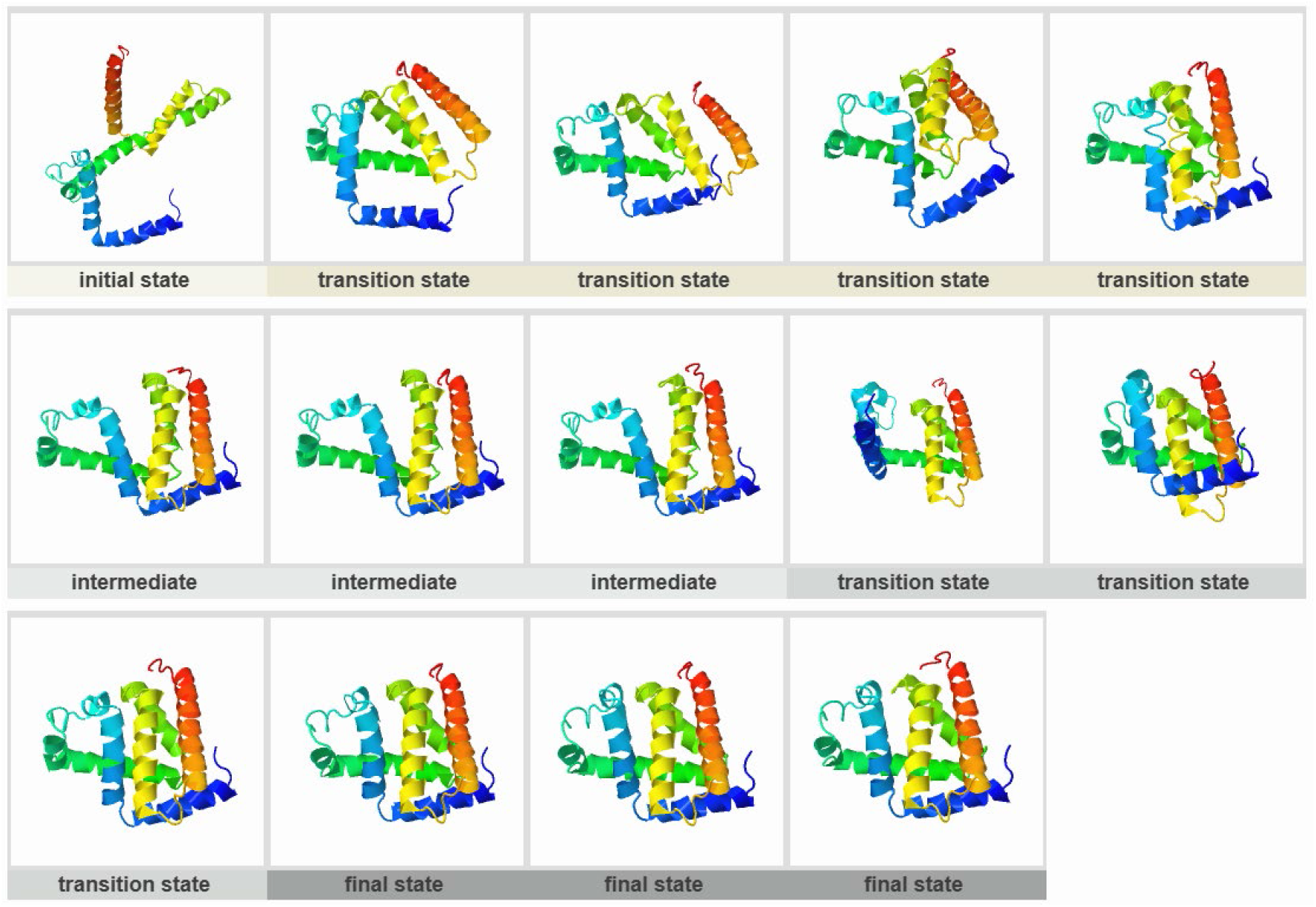
Apomyoglobin (PDB ID:1MBC)

**Figure S14.**
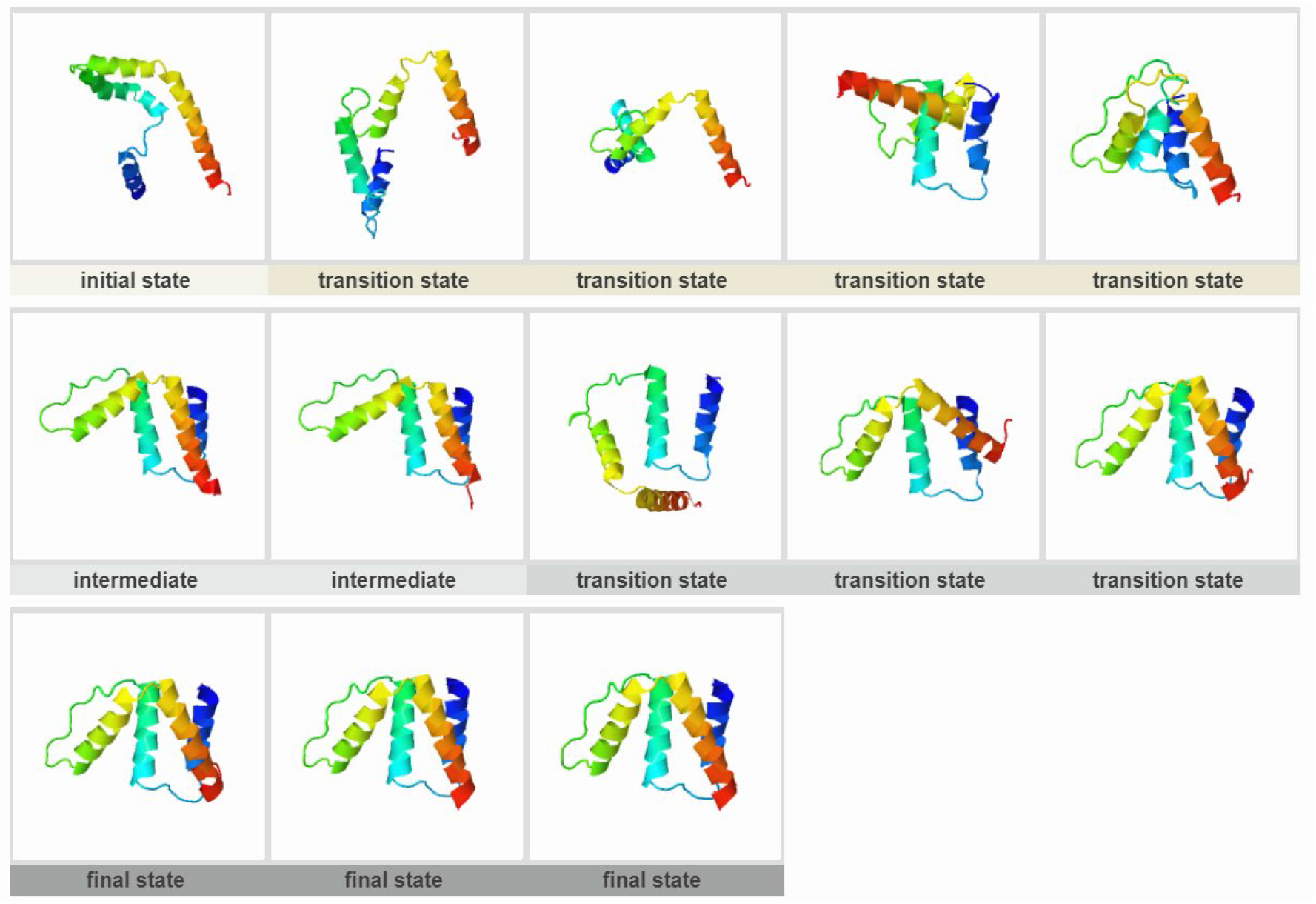
Acyl-CoA binding protein (PDB ID:1NTI)

**Figure S15.**
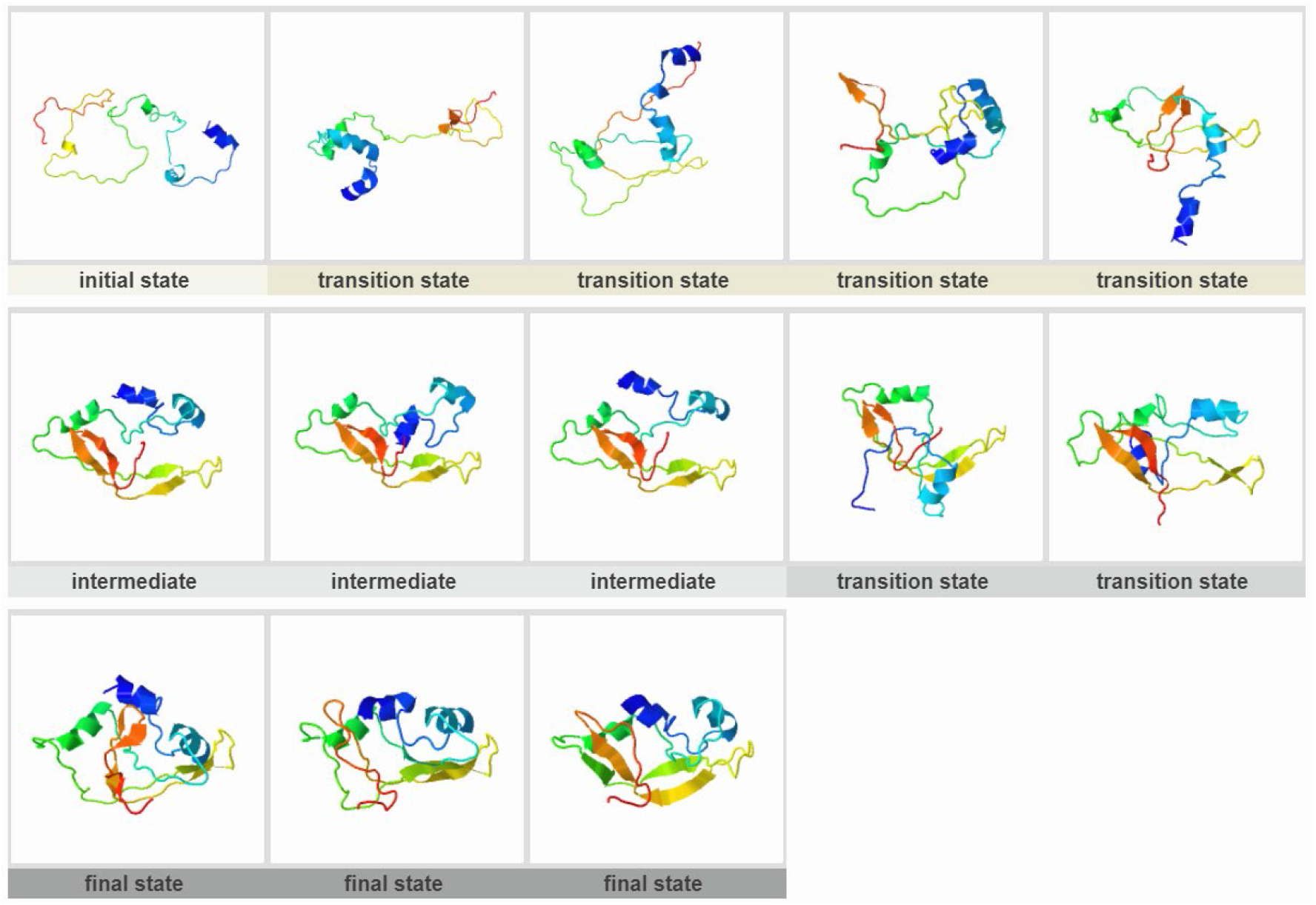
Onconase (PDB ID:1ONC)

**Figure S16.**
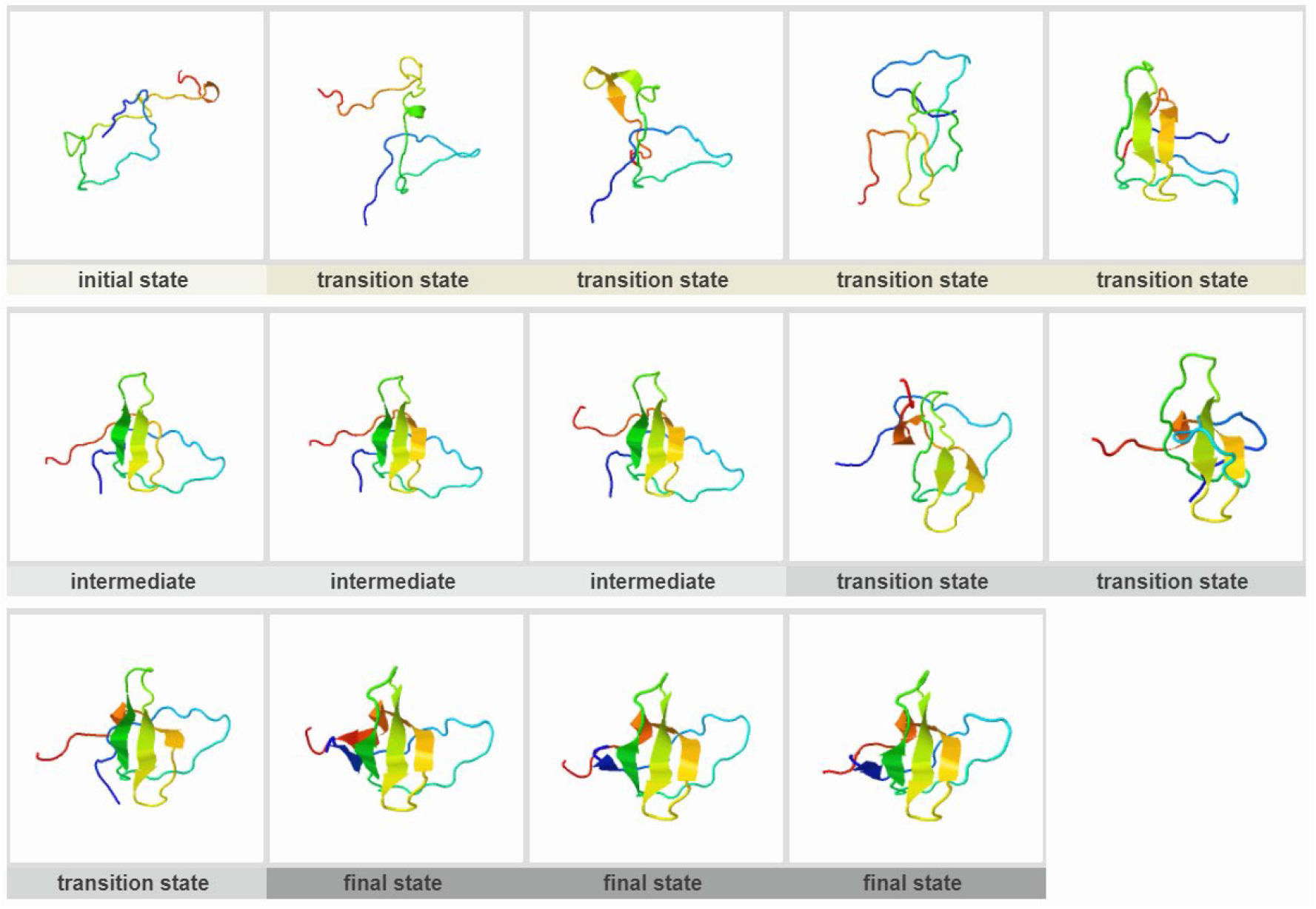
Fyn SH3 domain (PDB ID:1SHF)

**Figure S17.**
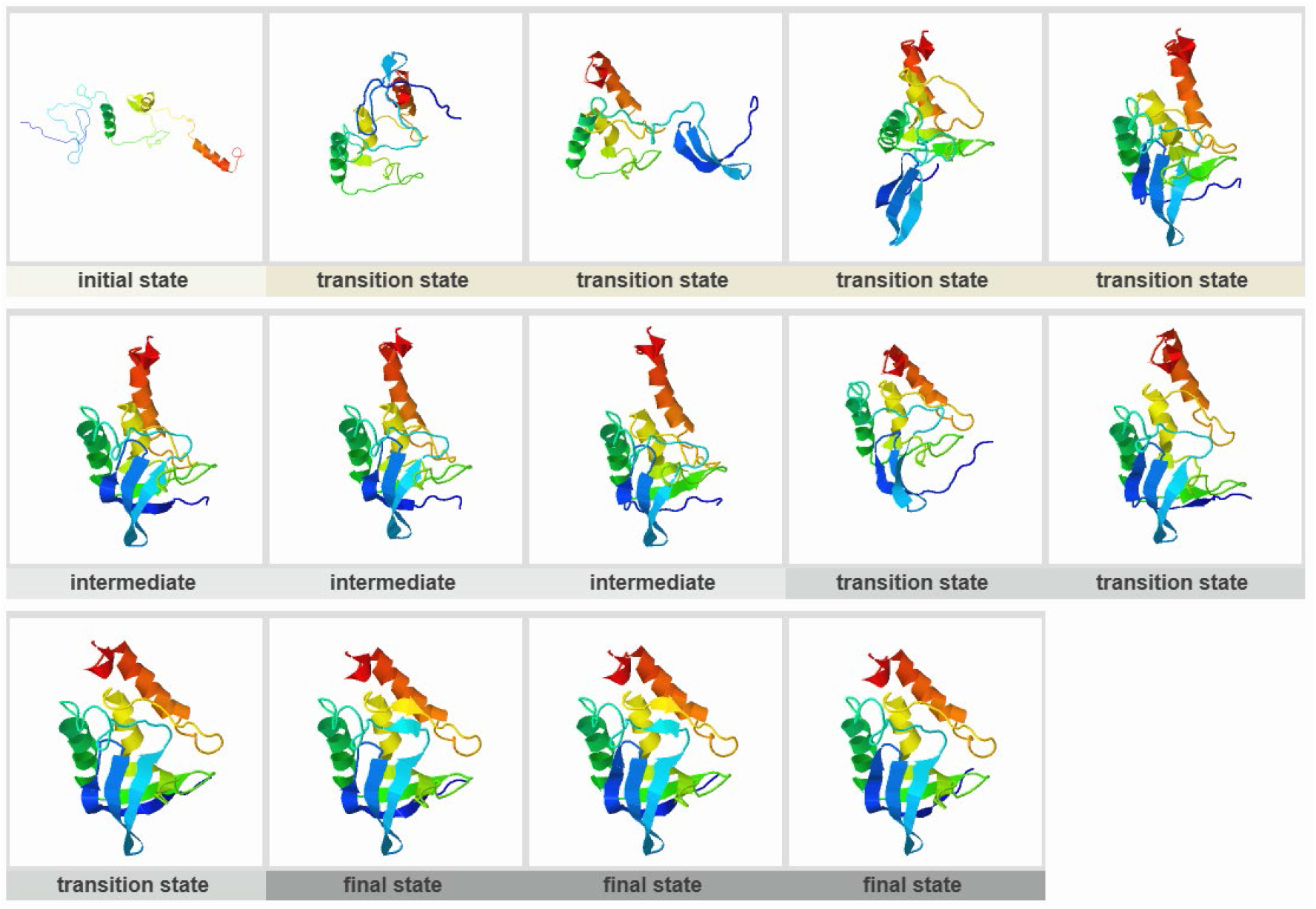
Staphylococcal nuclease (PDB ID:1STN)

**Figure S18.**
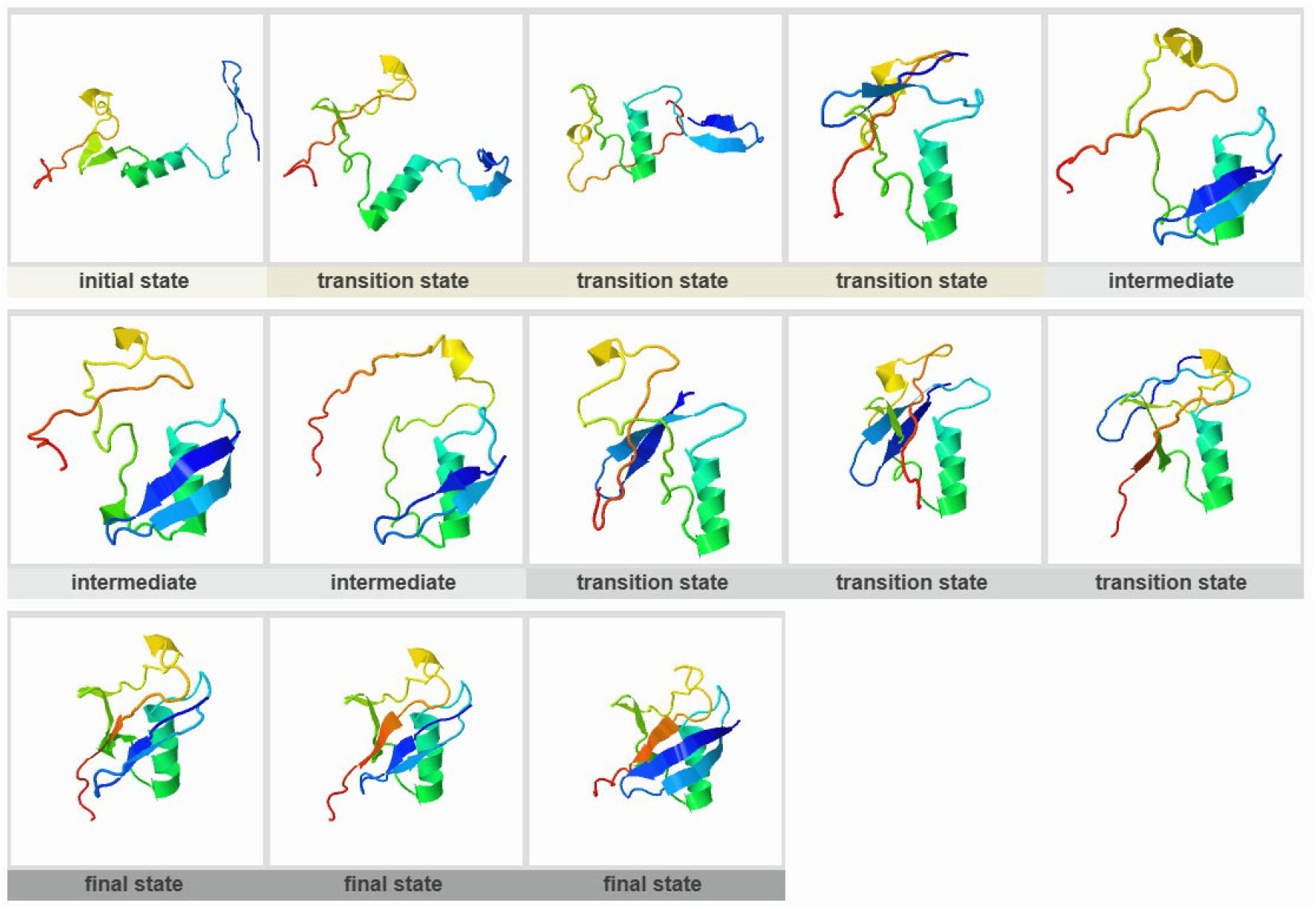
Polyubiquitin-C (PDB ID:1UBQ)

**Figure S19.**
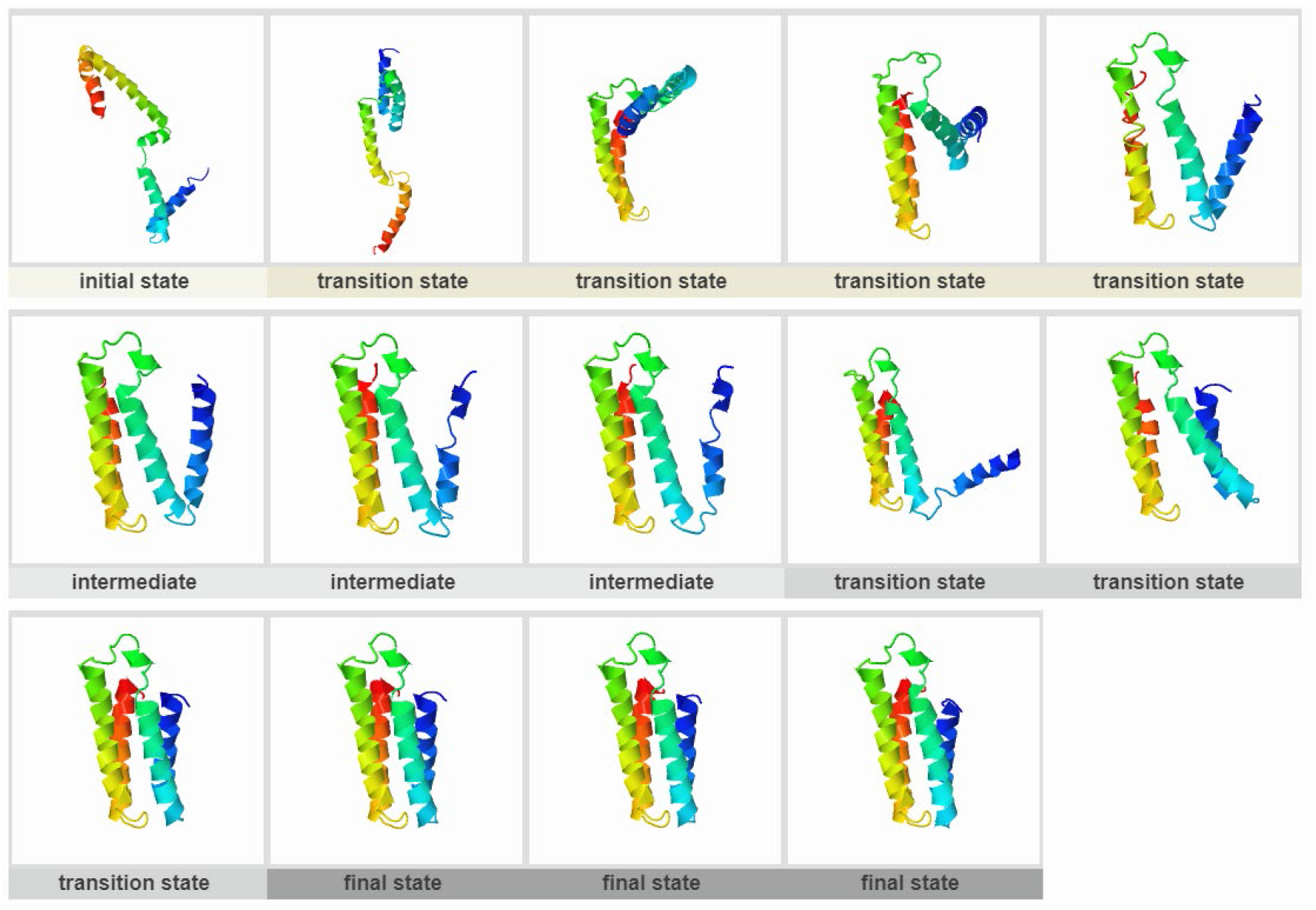
Rd-apocytochrome b562 (PDB ID:1YYJ)

**Figure S20.**
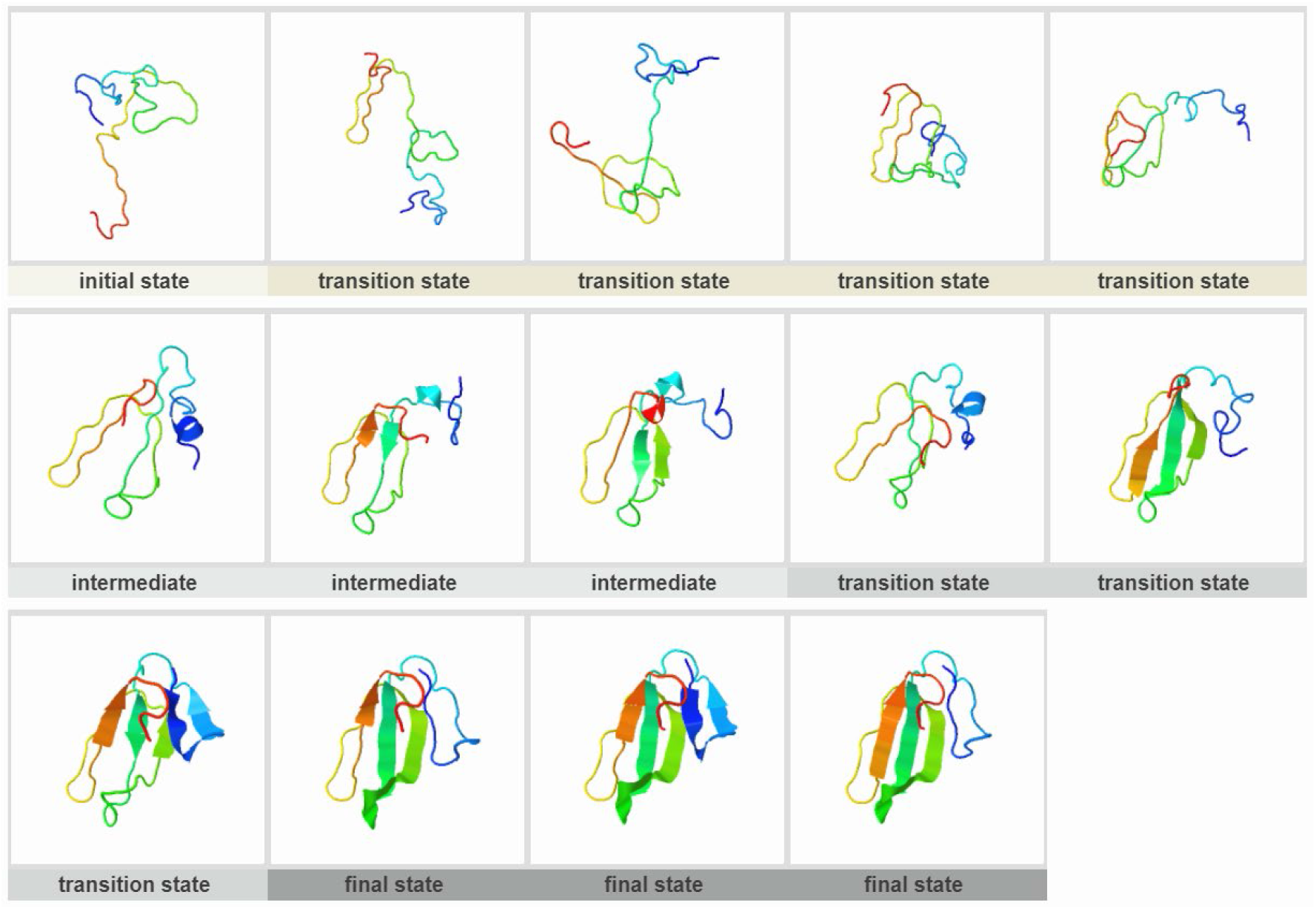
CTX III (PDB ID:2CRT)

**Figure S21.**
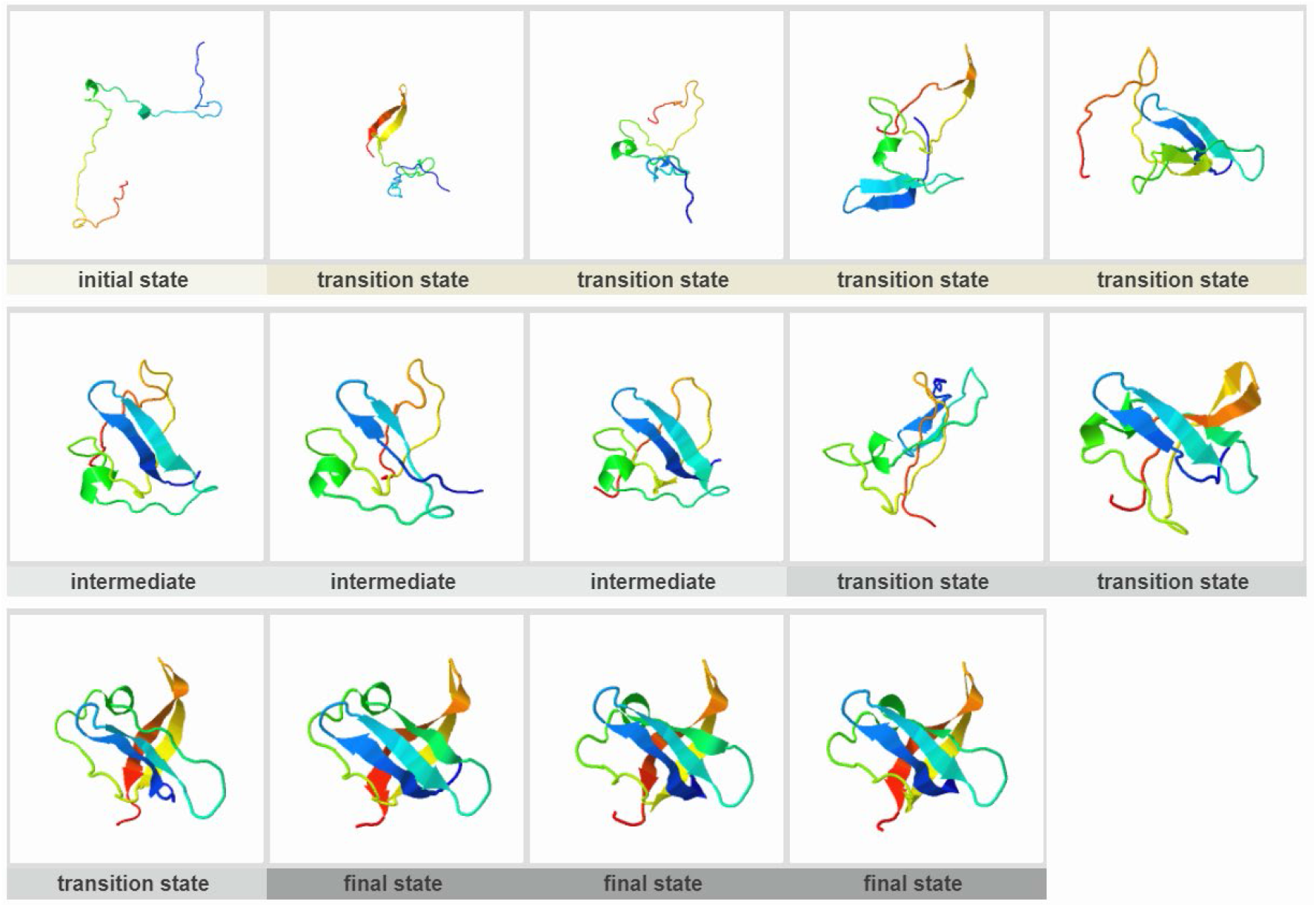
CspB (PDB ID:2F52)

**Figure S22.**
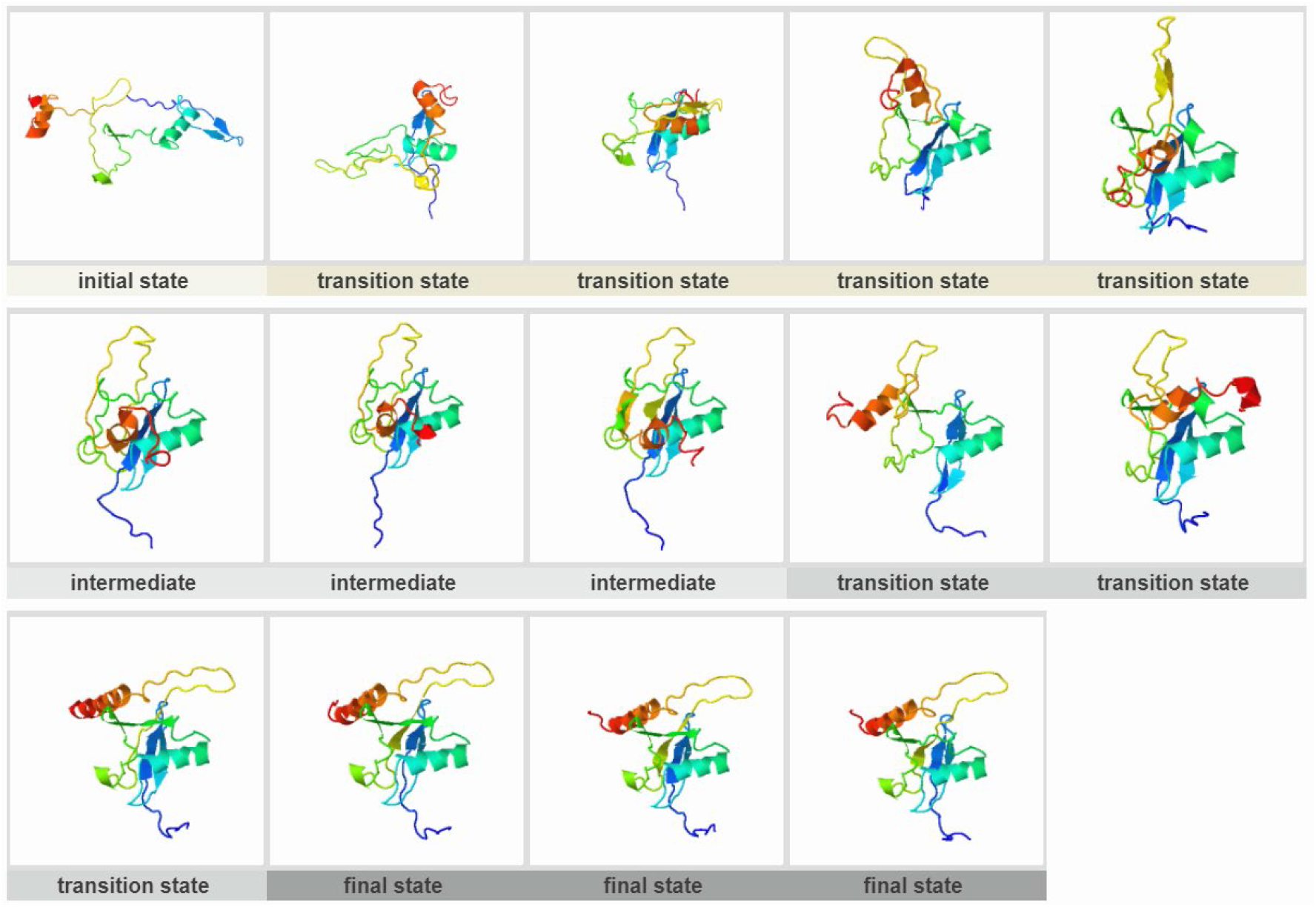
Ubq-UIM (PDB ID:2KDI)

**Figure S23.**
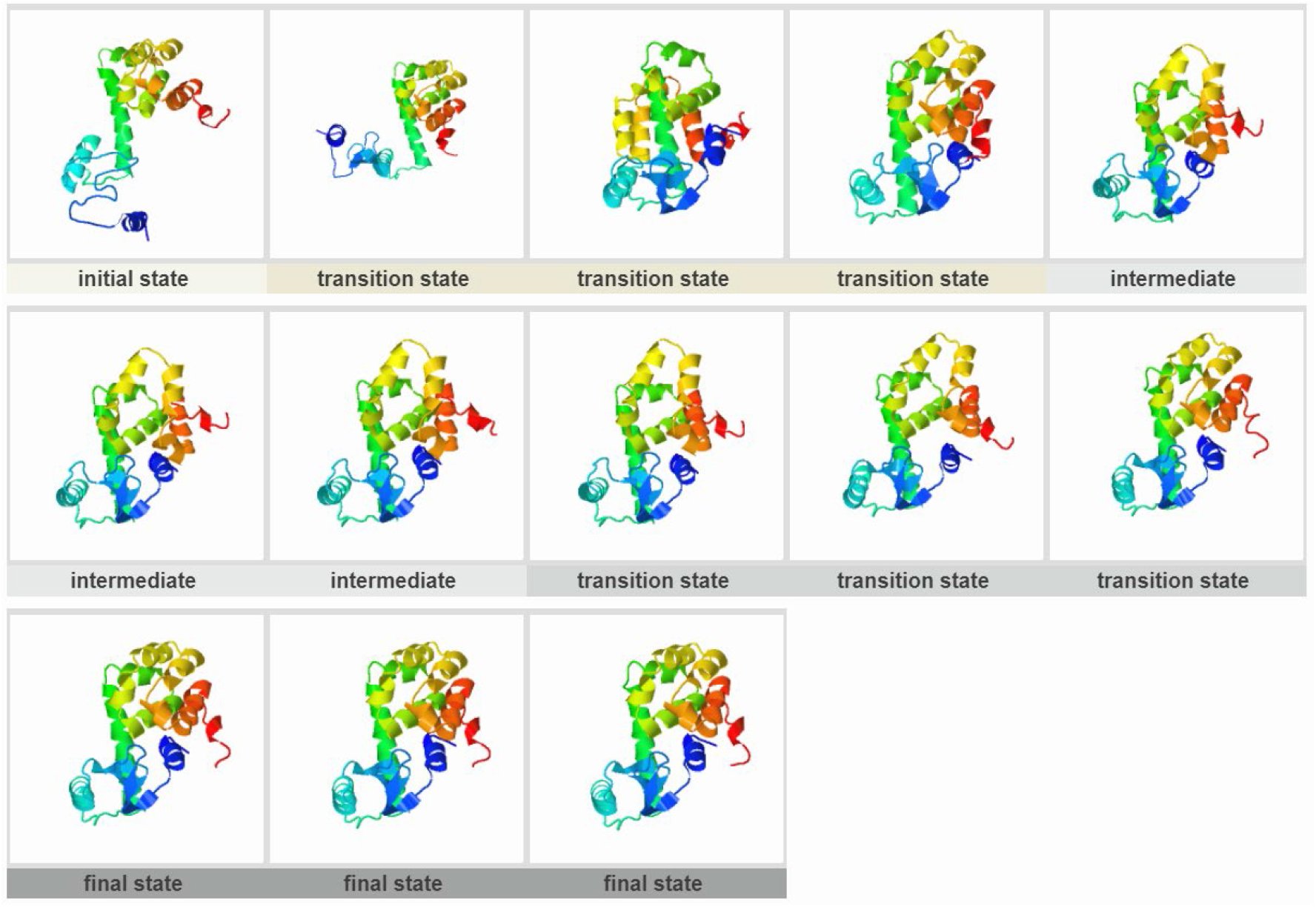
T4 Lysozyme (PDB ID:2LZM)

**Figure S24.**
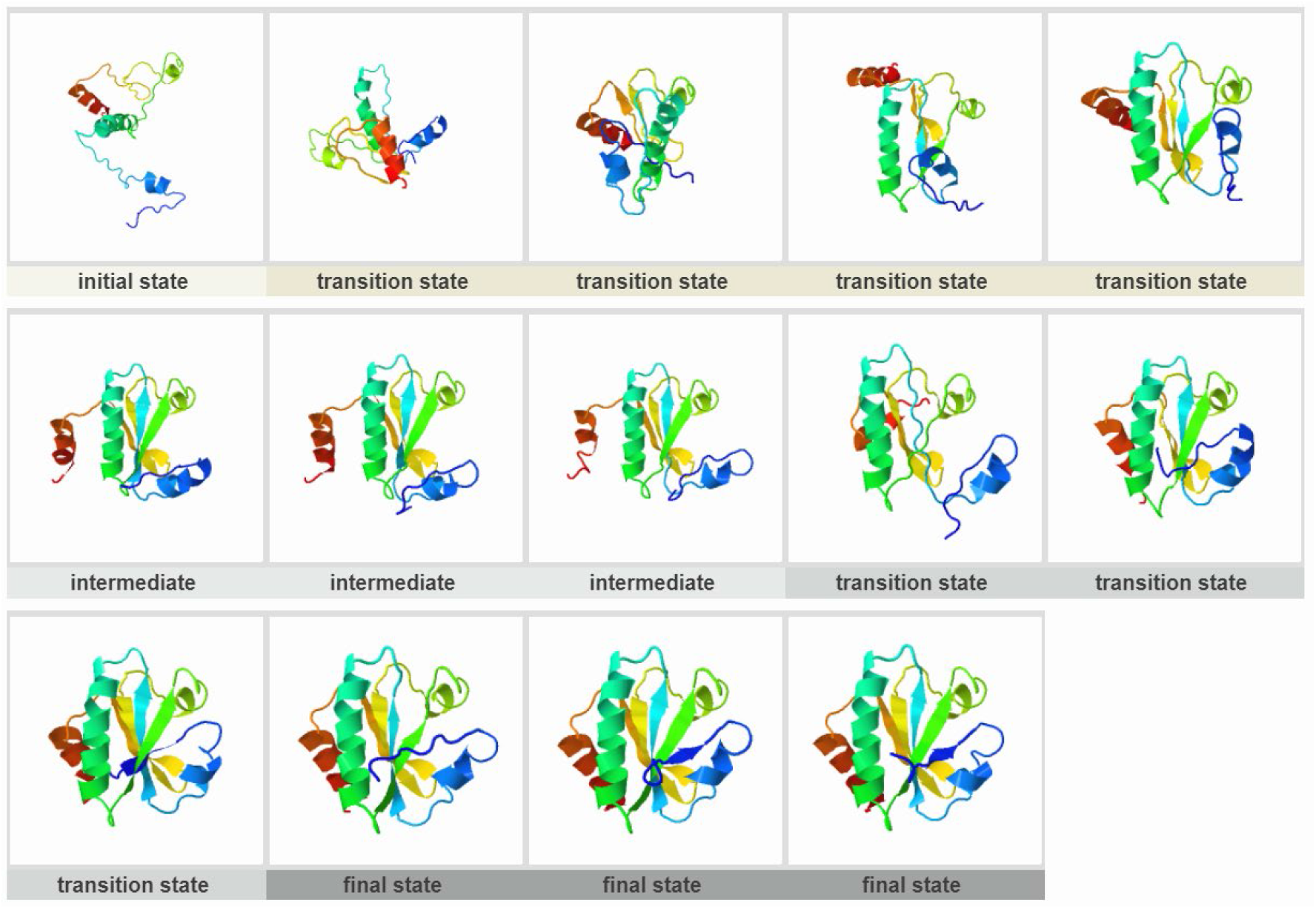
Thioredoxin (PDB ID:2TRX)

**Figure S25.**
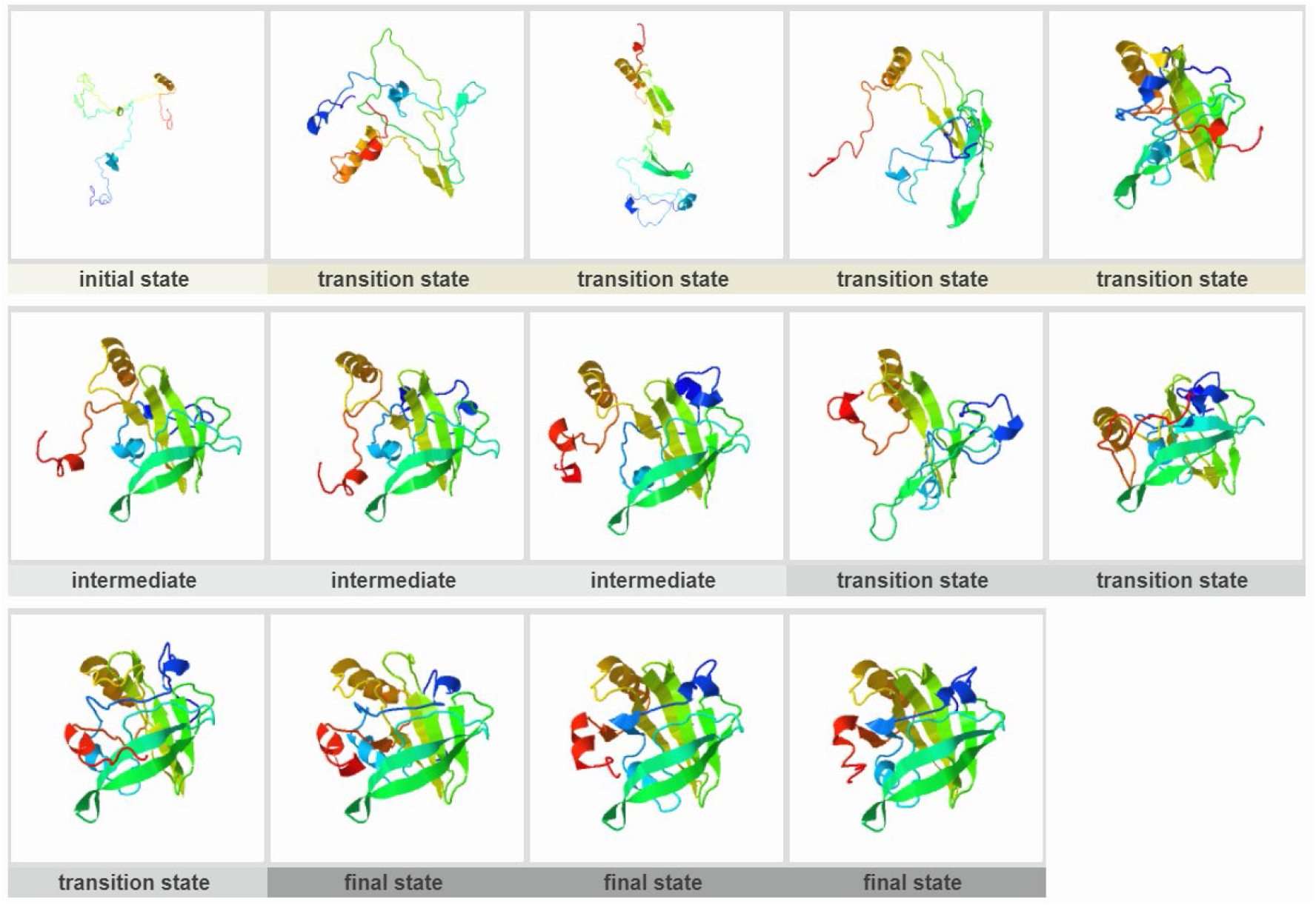
Beta-lactoglobulin (PDB ID:3BLG)

**Figure S26.**
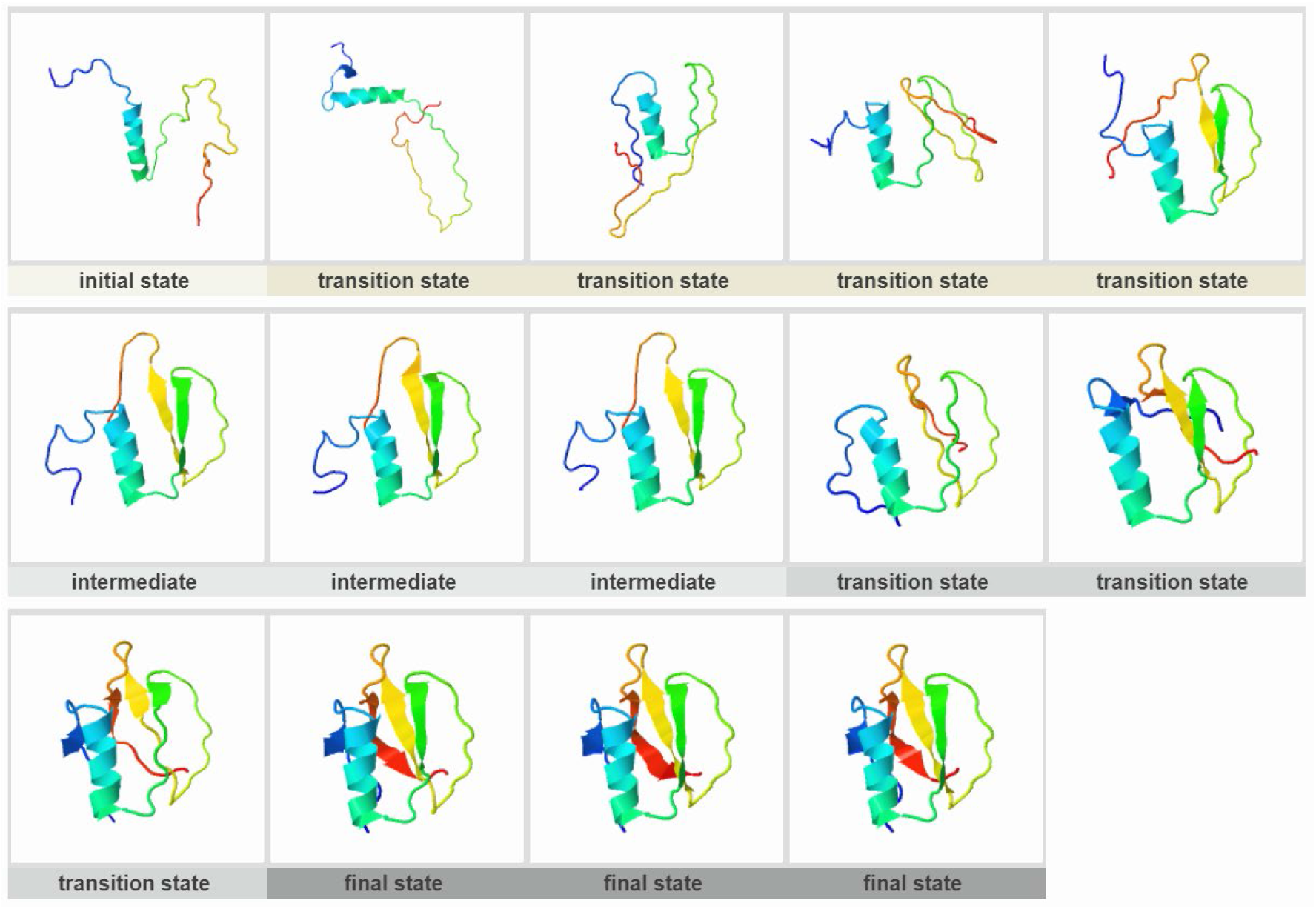
Chymotrypsin Inhibitor 2 (PDB ID:3CI2)

**Figure S27.**
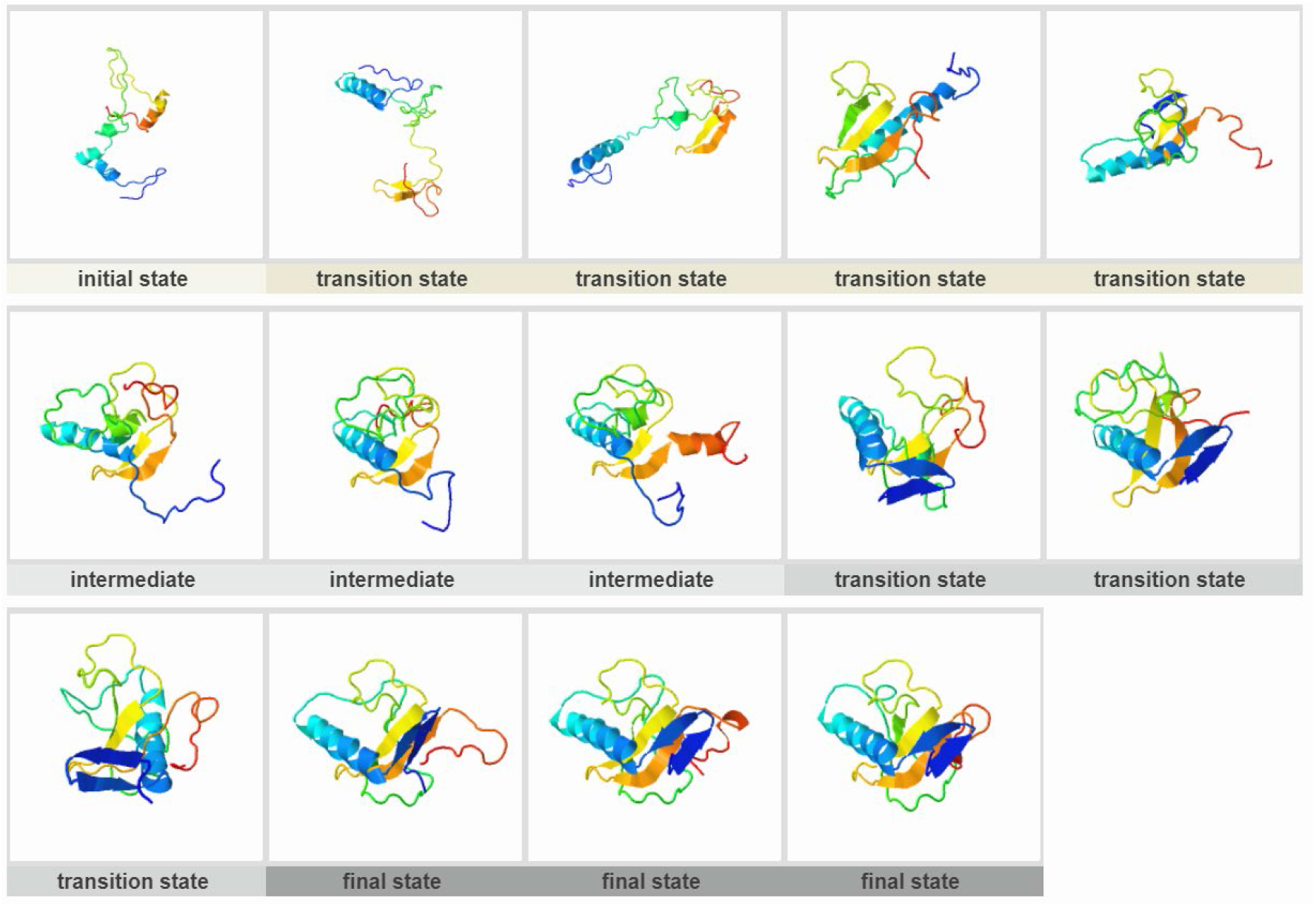
RNase T1 (PDB ID:3RNT)

**Figure S28.**
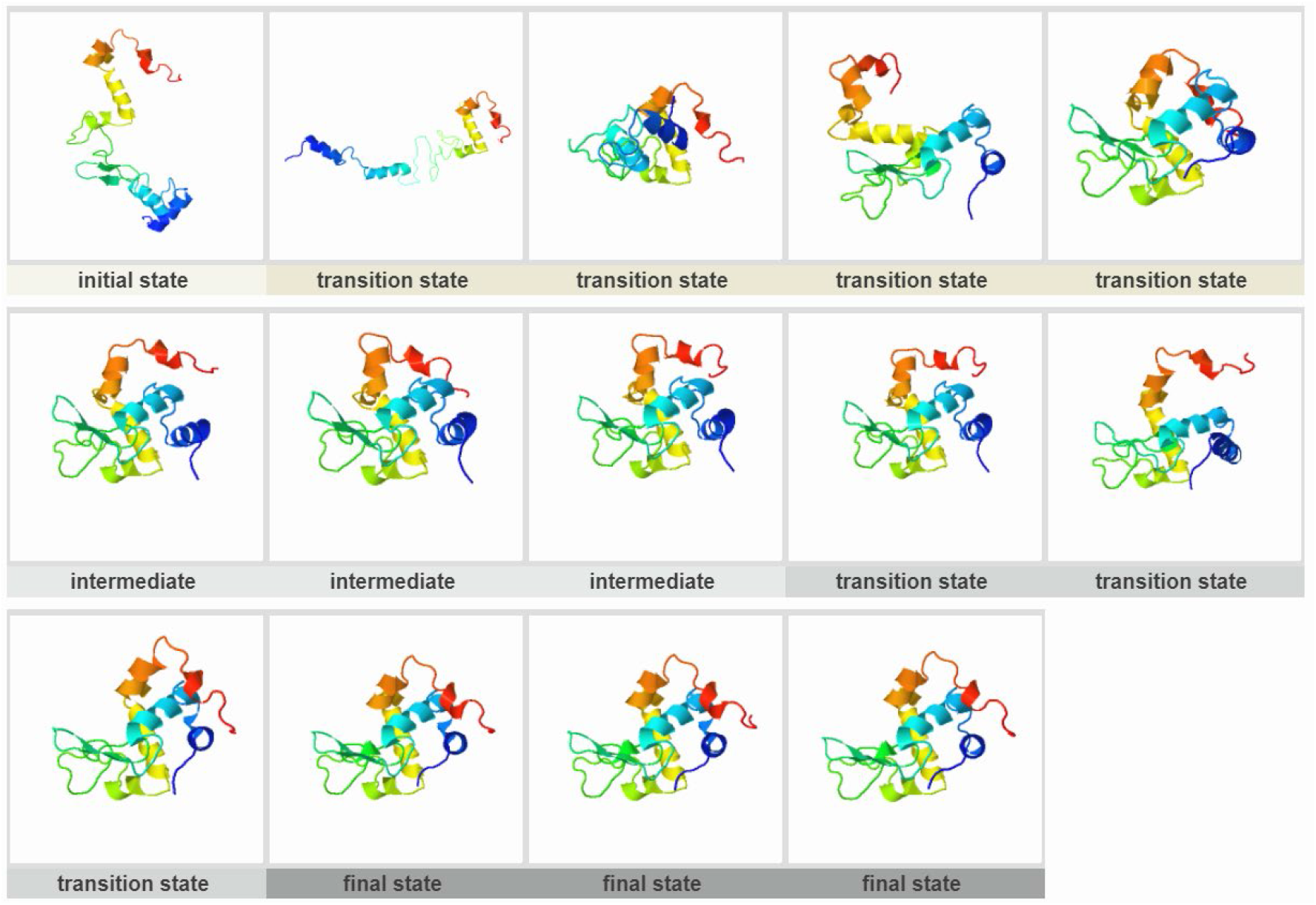
Lysozyme C (PDB ID:6LYZ)

**Figure S29.**
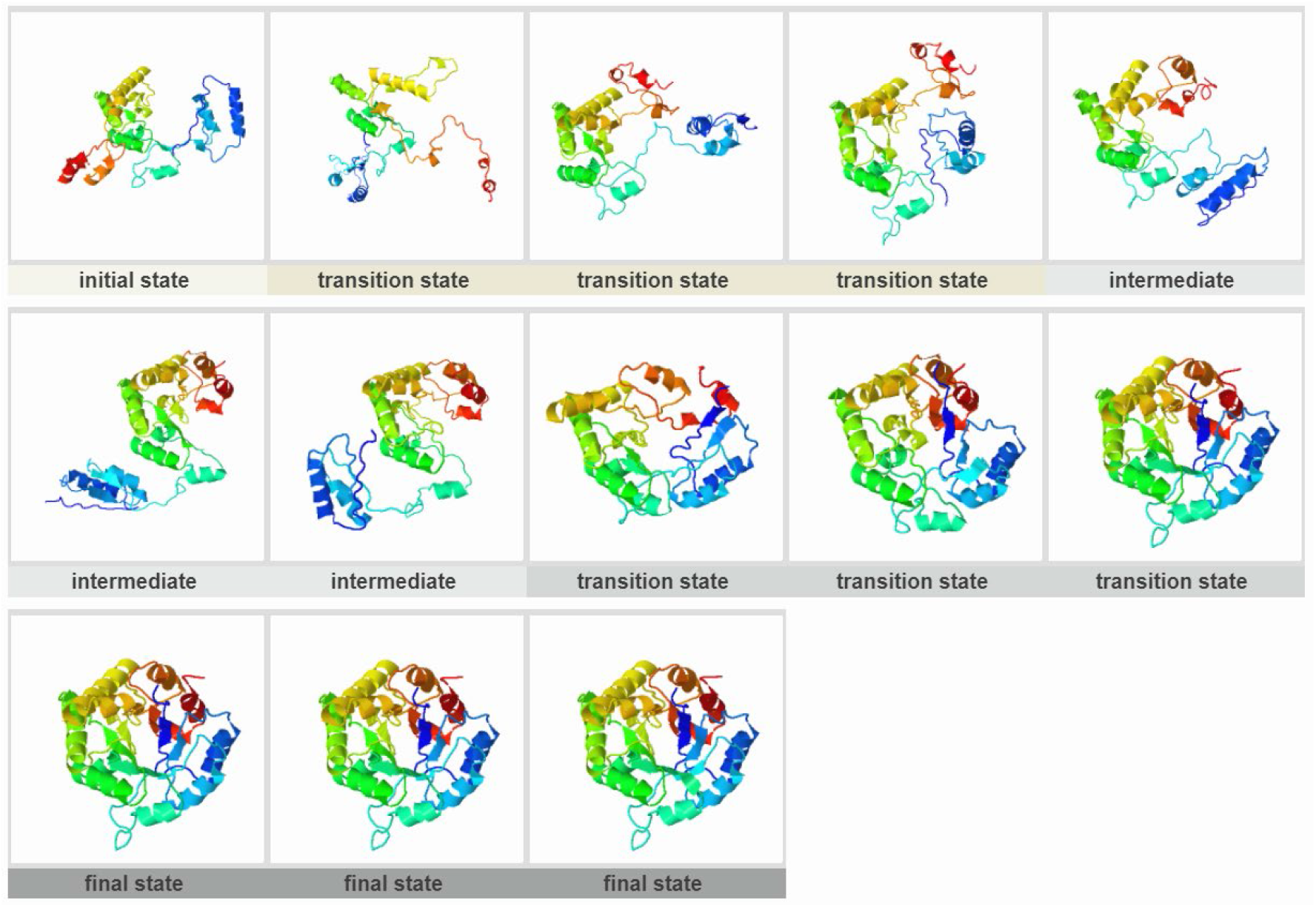
TIM (PDB ID:7TIM)

**Figure S30.**
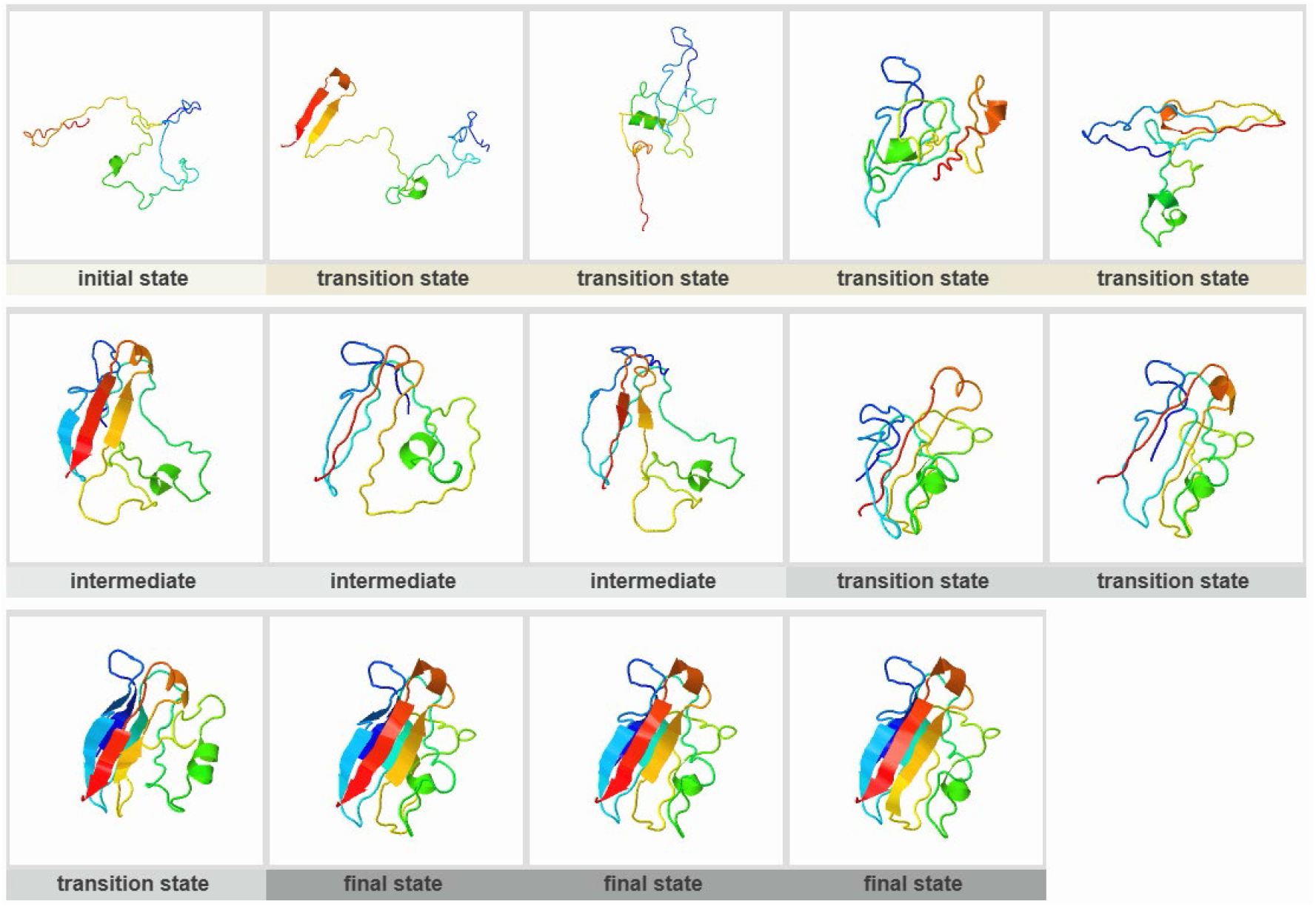
Plastocyanin (PDB ID:9PCY)

### Supporting Information Text

We collected descriptions and evidence of experimentally determined folding intermediates or transition states for 30 cases. They are present in (1) to (30).

#### (1) PDZ-3 domain

**Description from ref.[1]:** “Figure 1. The structure of PDZ-3 domain (pdb code: 1BE9). Residue F41 is mutated to Trp to provide a fluorescence probe. The hidden intermediate has the red region unfolded.”

“Figure 8. Proposed folding pathway for PDZ-3 with different concentrations of GdmCl. The blue filled oval represents the folded region of the intermediate. The red filled oval represents the region that is unfolded in the intermediate.”

#### (2) Barnase

**Description from ref.[2]:** “FIGURE 10: Free energy diagram illustration of the folding pathway of barnase under native conditions. Light gray represents unfolded regions. Colored regions represent folded regions.”

#### (3) BPTI

**Description from ref.[3]:** “In several of the folding trajectories, such as the one in figure, we find that in the conversion process between two native-like disulfide bond intermediates nonnative species, [30–51, 5–38] and [30–51, 5–14], are transiently populated. These were precisely the ones identified in experiments.”

#### (4) Apo-azurin

**Description from ref.[4]:** “The strongest consolidation of apo-azurin is observed in and around S3 with some diffuse contacts to S4–S6. Overall the structural consolidation is 43%.”

#### (5) Im7

**Description from ref.[5]:** “In the first millisecond of Im7 folding, a highly structured intermediate forms that contains three of the four native helices (Ⅰ, Ⅱand IV). The core of this intermediate is specific in that some of the hydrophobic side chains are not buried, most notably all of those that are highly exposed to solvent in the native state.”

#### (6) CTL9

**Description from ref.[4]:** “Overall this has one of the most weakly consolidated transition states with an average Φ_#_ of 0.21. However, a very clear clustering of some highly consolidated parts involving the S2-loop-S3 motif is observed.”

#### (7) Ckshs1

**Description from ref.[6]:** “The results show that ckshs1 folds sequential pairs of β-strands first (β1/β2 and β3/β4). Subsequently, these pairs pack against each other and onto the α-helical region to form the core.”

#### (8) FAS-associated death domain

**Description from ref.[7]:** “Thus, it appears that, while all six helices independently and cooperatively form concomitant with the hydrophobic collapse, only helices 1, 2, 4, and 5 interact in the transition state structure, with helices 3 and 6 associating on a later folding timescale.”

#### (9) Flavodoxin

**Description from ref.[8]:** “We suggest that the structured part of the putative intermediate is composed of the elements of secondary structure which have the slowest exchanging amide protons in the native protein. These elements are strands β1, β3, β4 and β5 and helices α4 and α5.”

#### (10) HIV-1 ribonuclease H

**Description from ref.[9]:** “Refolding of the isolated HIV RNase H domain shows a kinetic intermediate detectable by stopped-flow far UV circular dichroism and pulse-labeling H/D exchange. In this intermediate, strands 1, 4, and 5 as well as helices A and D appear to be structured.”

#### (11) Cytochrome c

**Description from ref.[10]:** “Experiment shows that, under equilibrium native conditions, cytochrome c unfolds by stepping energetically uphill through a ladder of forms that differ one from the next by the unfolding of one more native-like foldon (far right). HX MS experiments during kinetic folding demonstrate a pathway that steps sequentially downhill through the same intermediates. These results are able to specify the stepwise pathway in close to 3D structural detail (rather than as a 1D projection onto some reaction coordinate) because the downhill kinetic folding units and the uphill equilibrium unfolding units are very similar to the foldons that compose the native structure.”

#### (12) FKBP12

**Description from ref.[4]:** “The folding appears to have progressed already beyond initial nucleation. Strong interactions in the molecule are observed in and between many residues of S2, H, S5, S6.”

#### (13) Apomyoglobin

**Description from ref.[11]:** “Fig. 4. Location of the most rapidly protected amides. The above figure shows the location of the most rapidly protected amides. Distribution of fully and nonequilibrated amides in the 0.4-ms intermediate ensemble. The structure of holomyoglobin is represented as a tube of varying radius. Residues whose amides are fully equilibrated after 0.4 ms of refolding, i.e., for which [NH_open_ has reached the equilibrium value, are depicted in magenta with increased tube radius. Residues for which [NH_open_ has not reached the equilibrium values and which are fluctuating between folded and unfolded states are depicted by a thinner blue tube. Very thin tubes represent regions that do not exhibit exchange protection and probably remain unstructured in the folding intermediate.”

#### (14) Acyl-CoA binding protein

**Description from ref.[12]:** “The major folding transition state has been characterized in detail by value analyses of a large number of mutants, revealing that interactions between helices 1, 2, and 4 exist in this state.”

#### (15) Onconase

**Description from ref.[13]:** “On the other hand, I_Φ_ has well-established secondary structure involving the second helix and a large part of the α-sheet region, while the other parts of the protein remain unstructured in this intermediate.”

#### (16) Fyn SH3 domain

**Description from ref.[14]:** “The model’s folding intermediate is structured in the three β-strands that make up the protein’s core and is strikingly similar to intermediates detected in a recent NMR study of Fyn SH3 folding and to folding transition states elucidated in mutagenesis studies of SH3 domains. The unfolding intermediate is formed by dissociation of the folded protein’s two terminal β-strands from its core.”

“Transient intermediates on the protein’s folding pathways are represented by 50 structures collected from the simulation data as described in the text. The N-terminal (strand β1, RT loop), core (strands β2, β3, β4; n-src, distal loops), and C-terminal (3_10_ helix, strand β5) portions of the protein are drawn in blue, black, and red, respectively. Each of the structures in the ensembles is aligned with respect to strand β3.”

#### (17) Staphylococcal nuclease

**Description from ref.[15]:** “According to the previous NMR study, the N-terminal β1-β3 are the most stable structure formed at the first step of the folding process. Then, after the remaining C-terminal β4-β5 and α-helices are formed, the docking of the α-domain with the β-domain follows.”

#### (18) Polyubiquitin-C

**Description from ref.[16]:** “Most of the amide protons with a *P* value larger than 10^4^ are in the hydrophobic core formed by three strands of β-sheet and the α-helix, while those with *P* values less than 10^2^ are located in regions of irregular structure, or on the surface of the protein.” **Description from ref.[17]:** “Thus, the core of ubiquitin, composed of the a-helix and β-sheet and the interface between them, is formed in a major cooperative folding event. The sheet and helix protons are protected at nearly identical rates, indicating cooperativity in the formation both of the individual elements of secondary structure and of the interfacial region. The rapid protection rates for the amide protons of Ile-23 and Leu-56 provide direct evidence for early association of the sheet and the helix.”

#### (19) Rd-apocytochrome *b*_562_

**Description from ref.[18]:** “In the case of Rd-apocyt *b*_562_, a redesigned stable variant of apocytochrome *b*_562_ with a four-helix bundle fold, NHX experiments identified two partially unfolded forms (PUFs). PUF1 has the N-terminal helix and a part of the C-terminal helix unfolded. PUF2 has only the N-terminal helix unfolded.”

#### (20) CTX III

**Description from ref.[19]:** “Folding kinetics of CTX III based on the amide-protection data reveals that the triple-stranded, antiparallel β-sheet segment, which is located in the central core of the molecule, appears to fold faster than the double-stranded β-sheet segment.”

#### (21) cspB

**Description from ref.[20]:** “Thirdly, it highlights the residues in strand β4 (Phe49 and Ile51) and β5 (Ala60 and Val63) that are essential for the stability of the native protein (very high DDGNU values), but not of the transition state (low Φ-values). This suggests that the second β sheet is not stabilized in the transition state.”

“In the first β sheet (β1–β3) the Φ-values increase to 1 within the first half of strand β1, stay at 1 and then decrease to zero within the second half of strand β3. In the second sheet (β4–β5) the Φ-values are zero, and in the long connecting loop there is a narrow peak of intermediate to high Φ-values around residue 41. This led to the simple interpretation that in the transition state the first sheet is folded, but the second is unfolded.”

#### (22) Ubq-UIM

**Description from ref.[21]:** “We have previously engineered a Ubq-UIM fusion protein that allows independent experimental manipulations of the individual domains. It takes advantage of the fact that the ubiquitin interacting motif (UIM), consisting of a 20-residue helix, binds to ubiquitin (Ubq) in an orientation that positions the C-terminus of Ubq in close proximity to the N-terminus of the UIM.”

“Thermodynamic studies of cooperativity upon temperature-induced unfolding of Ubq-UIM showed that it does not follow two-state unfolding but includes detectable population of two equilibrium intermediate states consisting of either one of the two domains (Ubq or UIM) folded and the other unfolded.”

#### (23) T4 Lysozyme

**Description from ref.[22]:** “A native-state hydrogen exchange experiment subsequently detected an equilibrium intermediate that only involves the formation of the C-terminal domain.”

#### (24) Thioredoxin

**Description from ref.[23]:** “The experiments show that C-terminal α-helix is mainly unfolded in transition state ensembles TSE1 and the intermediate and becomes structured in TSE2. Structure-based molecular dynamics are in agreement with these experiments and provide protein-wide structural information on transient states. In our model, thioredoxin folding starts with structure formation in the β-sheet, while the protein helices coalesce later.”

#### (25) Beta-lactoglobulin

**Description from ref.[24]:** “The intermediate contains hydrogen bonded structure as measured by burst-phase labeling in the core of the β-sheet (βF, βG and βH) and the major α-helix, as well as some fluctuating helical structure near the N-terminus that is later converted into β-sheet.”

#### (26) Chymotrypsin Inhibitor 2

**Description from ref.[25]:** “Our results indicate a preferred pathway for the unfolding of Chymotrypsin Inhibitor 2: it starts with the loss of native contacts between N terminus and β3 and continues with β2 and β3. Some helical contacts and some non-native contacts around the termini persist even in highly unfolded conformations of the structure.”

#### (27) RNase T1

**Description from ref.[26]:** “All of the slowest exchanging amide residues are located in strands 2-4 of the central β-sheet and these residues are protected first in the early stages of folding. The residues that have somewhat lower rate constants for protection in early folding are predominantly found in the α-helix and the first strand of the small β-sheet.”

#### (28) Lysozyme C

**Description from ref.[27]:** “This species contains stable hydrogen bonded structure in the helical α-domain, but lacks stable structure in the second, predominantly β-sheet, domain. Formation of the α-domain intermediate occurs with biphasic kinetics.”

**Description from ref.[28]:** “The α-helical domain (residues 1-37 and 85-123) is shown in blue and the β-sheet domain (residues 38-84) in red.”

“The structure of the molten globule is highly heterogeneous, having the highly structured α-helical domain formed by loose hydrophobic interactions whereas the β-sheet domain is significantly more unfolded.”

#### (29) TIM

**Description from ref.[29]:** “To test the stability of the I_1A_ state, we further sampled another set of 100 trajectories beginning from the I_1A_ state for 8,000 time units. Even with a quadrupled simulation time, only 17% of the trajectories reached the native state, confirming the extremely long lifetime of the I_1A_ state.”

#### (30) Plastocyanin

**Description from ref.[30]:** “The amide protons protected in the intermediate are shown as white spheres attached to their amide nitrogen (dark spheres) with residue numbers.”

